# A humoral stress response protects *Drosophila* tissues from antimicrobial peptides

**DOI:** 10.1101/2023.07.24.550293

**Authors:** Samuel Rommelaere, Alexia Carboni, Juan F. Bada Juarez, Jean-Philippe Boquete, Luciano A. Abriata, Fernando Teixeira Pinto Meireles, Verena Rukes, Crystal Vincent, Shu Kondo, Marc S. Dionne, Matteo Dal Peraro, Chan Cao, Bruno Lemaitre

## Abstract

The immune response against an invading pathogen is generally associated with collateral tissue damage caused by the immune system itself. Consequently, several resilience mechanisms have evolved to attenuate the negative impacts of immune effectors. Antimicrobial peptides (AMPs) are small, cationic peptides that contribute to innate defenses by targeting negatively charged membranes of microbes^1, 2^. While being protective against pathogens, AMPs can be cytotoxic to host cells^1, 3^. Little is known of mechanisms that protect host tissues from AMP-induced immunopathology. Here, we reveal that a family of stress-induced proteins, the Turandots^4, 5^, protect *Drosophila* host tissues from AMPs, increasing resilience to stress. Deletion of several *Turandot* genes increases fly susceptibility to environmental stresses due to trachea apoptosis and poor oxygen supply. Tracheal cell membranes expose high levels of phosphatidylserine, a negatively charged phospholipid, sensitizing them to the action of AMPs. Turandots are secreted from the fat body upon stress and bind to tracheal cells to protect them against AMPs. *In vitro*, Turandot A binds to phosphatidylserine on membranes and inhibits the pore-forming activity of *Drosophila* and human AMPs on eukaryotic cells without affecting their microbicidal activity. Collectively, these data demonstrate that Turandot stress proteins mitigate AMP cytotoxicity to host tissues and therefore improve their efficacy. This provides a first example of a humoral mechanism used by animals limiting host-encoded AMP collateral damages.

## Introduction

Multiple mechanisms have evolved in animals to ensure homeostasis upon biotic and abiotic stresses. These include the production of heat-shock proteins, cryoprotectants, antioxidants, the unfolded protein response, and compensatory proliferation to replace dead cells, all of which contribute to resilience in stressful conditions^6–8^. These stress pathways also play an active role in defense against microbes, not only to prevent pathogen-induced damage but also to protect the host from deleterious effects of the immune response itself.

Antimicrobial peptides (AMPs) are small, cationic, usually amphipathic peptides that contribute to innate immune defense in plants and animals^1, 2, 9^. Most display potent antimicrobial activity *in vitro* by disrupting negatively charged microbial membranes. Host membranes of animals typically have a neutral charge and are therefore not affected by pore-forming AMP activity. However, AMPs can be cytotoxic to host cells in certain contexts when expressed at high levels^10–12^. Hypotheses suggest that AMPs could target cell membranes that become negatively charged due to translocation of the phospholipid phosphatidylserine (PS) to the outer leaflet upon stress^13, 14^. To date, little is known of host mechanisms that protect tissues against AMP collateral damage during the immune response.

Antimicrobial peptides are well characterized in the fruit fly *Drosophila melanogaster* where they enable resistance of microbial infections and shape the microbiota^1, 15, 16^. In this insect, systemic infection triggers massive secretion of multiple AMPs by the fat body, an equivalent of the mammalian liver. AMPs can also be induced by abiotic stressors such as osmotic stress or desiccation^17, 18^. In *Drosophila*, the *Turandot* (*Tot*) gene family produces eight small secreted proteins that are highly expressed in AMP-like patterns during stress and immune responses^4, 5^. Although expression of *Tot* genes is widely used as a readout of the stress response^19–21^, their molecular function is unknown. Here we show that Turandot proteins are humoral factors that protect host tissues, notably the respiratory epithelium, from antimicrobial peptides, contributing to stress resilience and host defense.

## Results

### Turandot-deficient flies have low resilience to stress

Because of their sequence similarity and overlapping expression patterns, we anticipated functional redundancies among the eight *Turandot* genes. To assess their function, we generated compound mutants inactivating up to 6 out of 8 Turandot genes. We first generated a mutant deleted for a genomic cluster of four genes (*TotA*, *TotB*, *TotC* and *TotZ*) called *Tot^AZ^*. In this mutant, we then inactivated *TotM*, *TotX,* or both, creating the *Tot^MAZ^*, *Tot^AZX^* or *Tot^XMAZ^* lines, respectively. Tot-deficient flies were viable and fertile. The expression pattern of *Turandot* genes implied that they might play an important role against a broad range of stresses^4, 19, 20, 22, 23^. We therefore subjected *Tot*-deficient flies to biotic and abiotic challenges. *Tot^XMAZ^* animals showed a mild, yet not significant, susceptibility to systemic infection with *Drosophila* C virus, *Pectobacterium carotovorum carotovorum* (*Ecc15*) or *Enterococcus faecalis* (**Extended Data Fig. 1a-c**), as compared to isogenic wild-type flies. Strikingly, *Tot^XMAZ^*flies were more susceptible than wild-type animals to starvation, heat, and osmotic stresses (**Fig. 1A-C**). *Tot^AZ^* mutants showed a mild and variable susceptibility to these stresses, while both *Tot^MAZ^* and *Tot^AZX^* displayed a strong and consistent susceptibility to these challenges. *Tot^AZX^* and*Tot^XMAZ^* flies were similarly susceptible to all challenges tested (**Fig. 1D**). Thus, we mostly used *Tot^AZX^* and *Tot^XMAZ^* flies to address Turandot function in the following genetic characterization (see methods). Overexpression of *TotA* alone in the fat body of *Tot^XMAZ^* flies partially rescued resistance to osmotic stress (**Fig. 1E**), confirming that the susceptibility is caused by *Tot* deficiency. Interestingly, ubiquitous overexpression of *TotA* in the *Tot^XMAZ^*background failed to rescue the survival phenotype, suggesting that ectopic or excessive expression of *TotA* is detrimental to flies (**Extended Data Fig. 1d,e**). Together, these data demonstrate that *Turandot* genes are required for optimal resilience to a broad set of environmental challenges.

**Figure 1:**
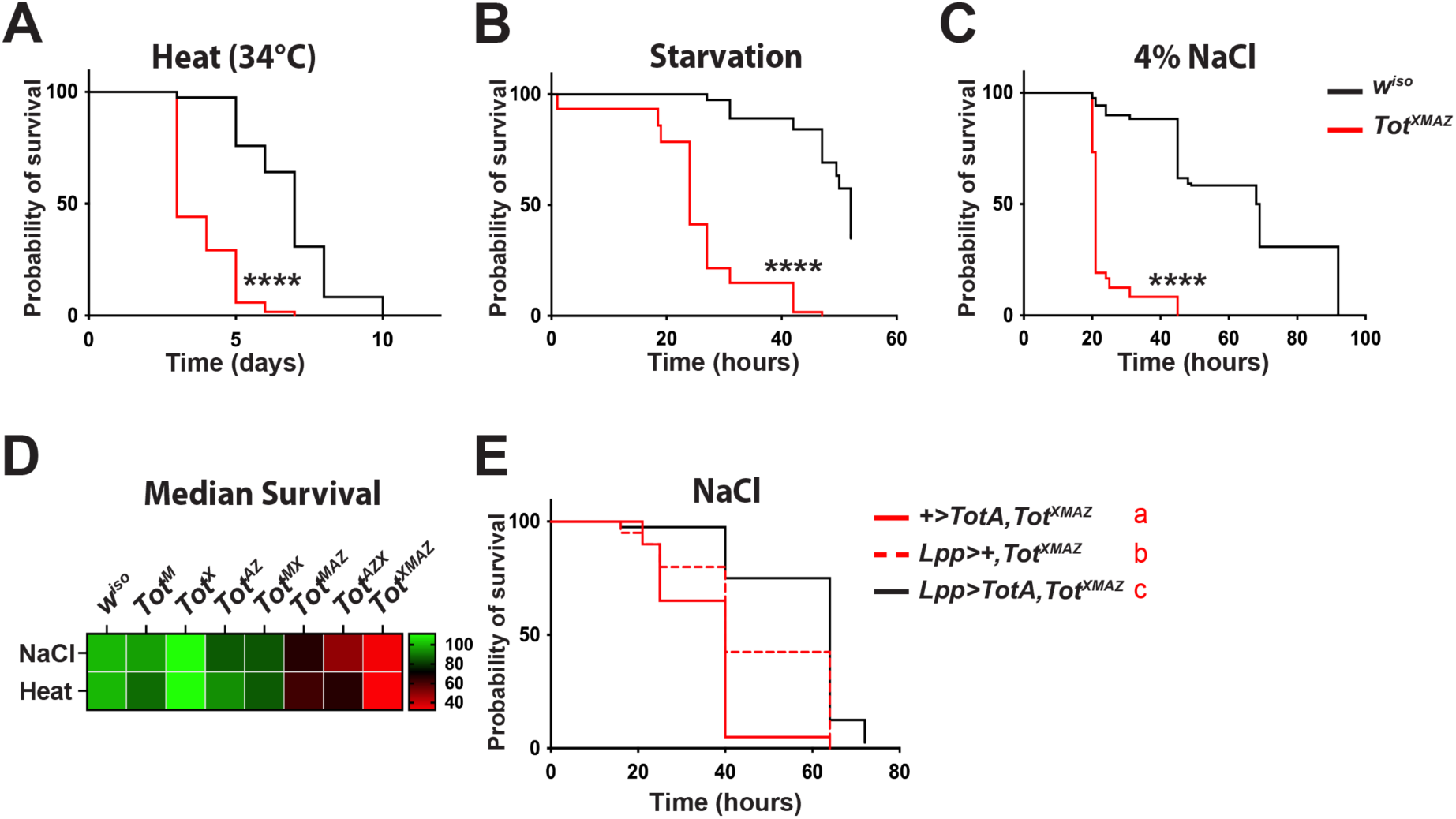
Turandot-deficient flies are susceptible to stress. **A-C**. Survival of wild-type *w^iso^*control (black, n=120) and *Tot^XMAZ^* flies (red, n=120) maintained at 34°C (**A**) or on agar only (**B**) or on food containing 4% NaCl (**C**). **D**. Heatmap representing the median survival of different *Tot*-deficient fly lines to heat and osmotic stresses (normalized to *w^iso^*). **E**. Survival upon osmotic stress of *+>TotA;Tot^XMAZ^* (red, solid line, n=60), *lpp>+;TotA;Tot^XMAZ^* (red, dashed line, n=80) or *Tot^XMAZ^* overexpressing *TotA* in the fat body (*lpp>TotA;Tot^XMAZ^*black, n=80). Results shown are a pool of 3 independent experiments. (****, *p*<0.00005; CoxPH test, followed by bonferonni correction when applicable. In this case, a compact letter display was used to show the statistics (see methods)).

### Secreted TotA binds to trachea

We next analyzed the expression pattern of Tot proteins, focusing on TotA. Western blot with an anti-TotA polyclonal antibody confirmed secretion of this protein into the hemolymph (insect circulatory fluid) (**Fig. 2A**). Immunostaining also revealed the presence of TotA in the fat body and, to a lesser extent, on trachea (insect respiratory system) and visceral muscles of unchallenged flies (**Fig. 2B and Extended Data Fig. 2a,b**). TotA was mostly localized at the plasma membrane and in intracellular punctae **(Fig. 2B)**. Osmotic stress and bacterial infection increased TotA staining in the fat body and the trachea (**Fig. 2C and 2D**). Because TotA is weakly expressed in trachea in basal conditions (**Extended Data Fig. 2c**), we hypothesized that this staining could result from tracheal binding of TotA secreted by the fat body. Consistent with this, HA-tagged TotA was observed on trachea when remotely expressed in the fat body **(Fig. 2E)**. Similar results were obtained with TotM overexpression, suggesting that fat body-derived Tots bind to trachea (**Extended Data Fig. 2d**). Conversely, knocking down TotA in the fat body reduced TotA staining of the trachea (**Fig. 2F**). Finally, we injected recombinant TotA protein into *TotA*-deficient animals and detected it on the plasma membranes of the fat body and trachea (**Fig. 2G**). These data show that upon stress TotA is secreted into the hemolymph and binds to fat and tracheal cells.

**Figure 2:**
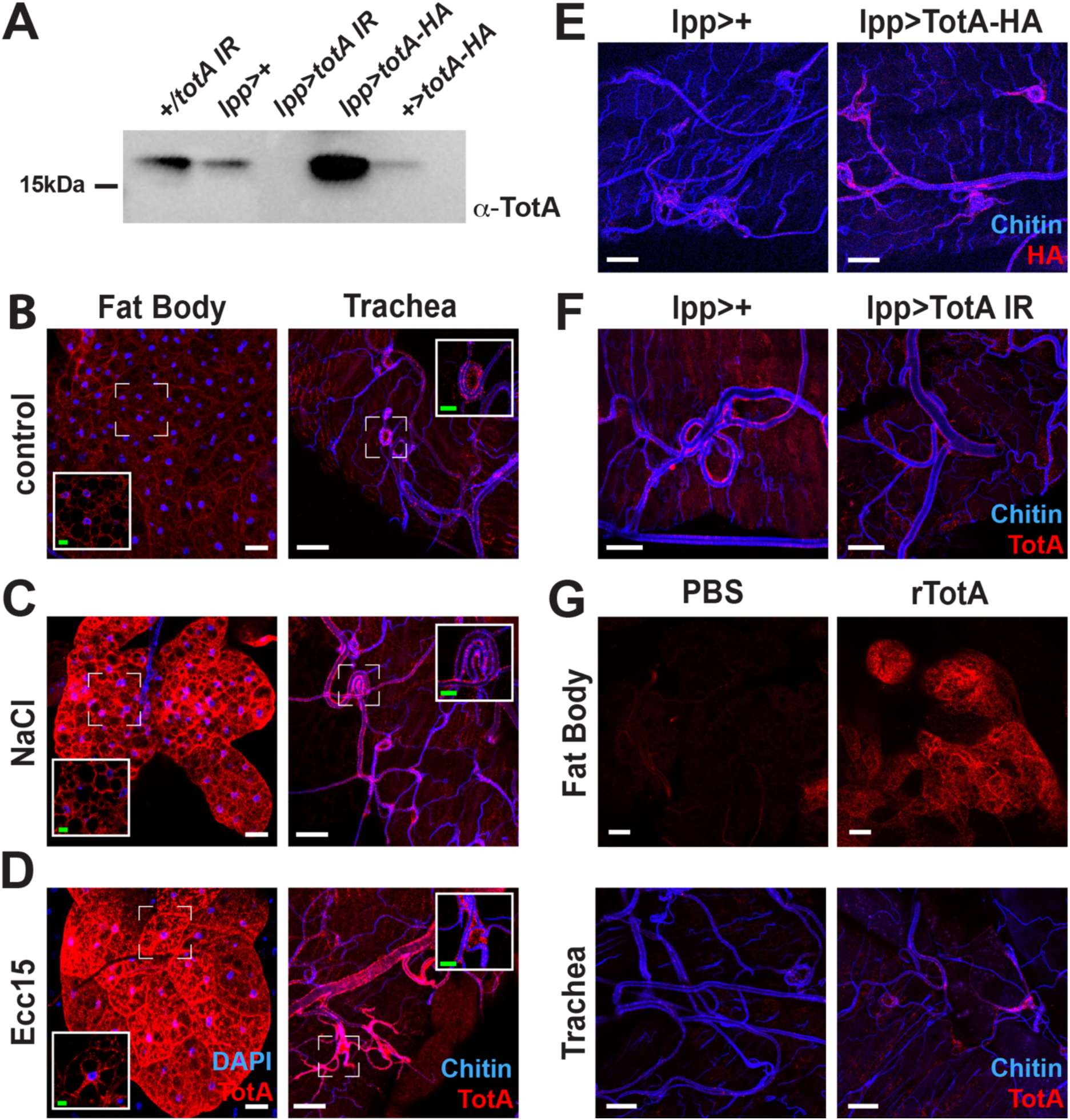
Fat body-secreted Turandots bind to trachea. **A**. Anti-TotA Western blot analysis of hemolymph from animals overexpressing TotA-HA or TotA RNAi in the fat body. **B-D**. Anti-TotA immunostaining of fat bodies (left) or gut trachea (right) of unchallenged flies (**B**) or flies exposed to osmotic stress (NaCl, **C**) or *Ecc15* infection (**D**). Insets show high magnification of the area defined by the white dotted square. Image brightness in these insets was adjusted to highlight the punctated structures **E**. Anti-HA immunostaining of trachea from control (left) or TotA-HA overexpressing (right) flies. **F**. Anti-TotA immunostainings of trachea from wild-type flies (left) or flies overexpressing TotA RNAi in the fat body (right). **G**. Anti-TotA immunostaining of fat bodies (upper panels) and trachea (lower panels) from flies injected with PBS (left) or with recombinant TotA. Blue, chitin; Red, TotA or TotA-HA. Scale bar: white, 20μm, green, 5 μm.

### Turandots maintain tissue oxygenation by protecting trachea from apoptosis

The binding of fat body-produced Tot proteins to trachea was unexpected. Therefore, we further investigated the role of Turandots in the respiratory epithelium upon stress, focusing on the tracheal network around the gut. We found that *Tot^XMAZ^* mutant flies had fewer terminal tracheal cells (TTC), and reduced tracheal coverage of the midgut compared to wild-type flies, as revealed by autofluorescence^24^ (**Fig. 3A-C**) or a *dSRF>GFP* tracheal reporter^25^ (**Fig. 3D**). Overexpressing *TotA* in the fat body of *Tot^XMAZ^* flies was sufficient to restore TTC numbers comparable to wild-type flies (**Fig. 3E)**. As we did not detect any anatomical defect in the trachea of *Tot^XMAZ^*larvae (**Extended Data Fig. 3**), we speculated that Turandots were required during metamorphosis to ensure proper tracheal morphogenesis, consistent with the high expression of Tots at the pupal stage^26^. Adult tracheal branching is modulated by nutrient cues and infection^24, 27, 28^, which prompted us to test whether stress could affect tracheation. Osmotic stress significantly reduced TTC number and tracheal coverage in wild-type animals to the level observed in unchallenged *Tot^XMAZ^* flies, showing that the tracheal system is intrinsically vulnerable to environmental stresses (**Fig. 3B and 3C**). Osmotic stress did not further reduce TTC number in *Tot^XMAZ^* flies, suggesting that tracheation was reduced in these flies regardless of challenge.

**Figure 3:**
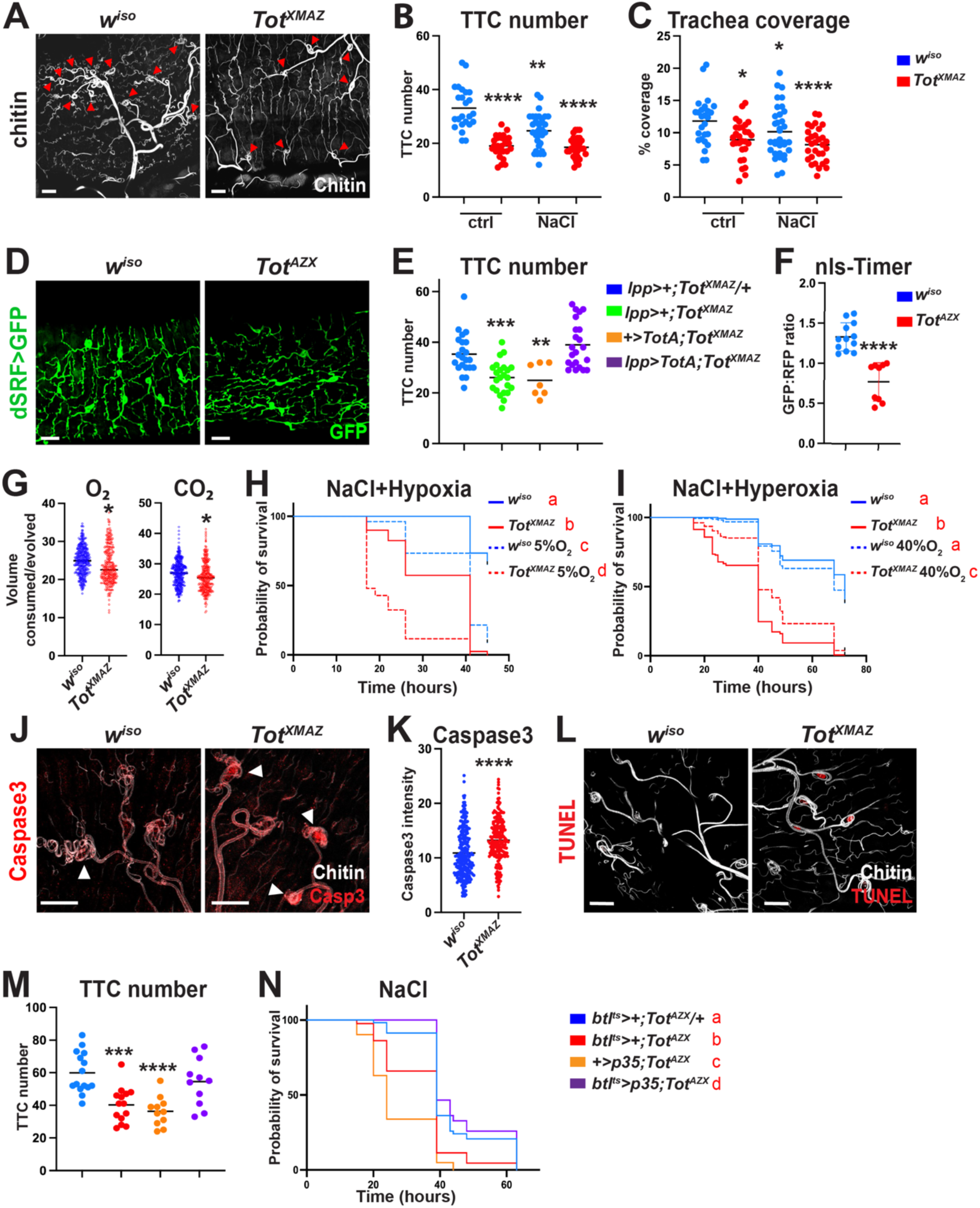
Turandots promote resilience to stress by preventing tracheal apoptosis. **A**. Chitin autofluorescence (grey) of gut trachea of *w^iso^* (left) or *Tot^XMAZ^* (right) flies. Red arrowheads indicate TTC. **B,C**. Quantification of gut TTC (**B**) and tracheal coverage (**C**) from *w^iso^* (blue) and *Tot^XMAZ^* (red) flies in unchallenged condition or exposed to osmotic stress. **D**. Anti-GFP immunostaining (green) of gut trachea from *dsrf>GFP* (left) or *dsrf>GFP,Tot^AZX^* (right) flies. **E**. Expression of TotA by the fat body Gal4 driver (*lpp-Gal4*) restores wild-type TTC counts in *Tot^XMAZ^* flies. **F**. Ratiometric analysis of the oxygen *nls-timer* fluorescence in wild-type (blue) or *Tot^AZX^* flies (red). **G**. O_2_ consumption (left) and CO_2_ production (right) of *w^iso^* (blue) or *Tot^XMAZ^* (red) flies. **H,I**. Survival to osmotic stress of *w^iso^* (blue) and *Tot^XMAZ^* (red) flies kept in normoxia (solid line), hypoxia (dashed line, **H**) or hyperoxia (dashed line, **I**). **J**. Cleaved caspase-3 (red, arrowheads) staining of trachea (chitin, white) from *w^iso^* (left) or *Tot^XMAZ^* flies exposed to osmotic stress. **K**. Quantification of caspase-3 staining intensity in TTC from *w^iso^* (blue) or *Tot^XMAZ^* (red) flies fed NaCl food. **L**. TUNEL staining (red) staining of trachea (chitin, white) from *w^iso^* (left) or *Tot^XMAZ^* flies exposed to osmotic stress. **M,N.** Gut TTC quantification (**M**) and survival to osmotic stress (**N**) of *Tot^AZX^* flies overexpressing p35 in trachea using the *btl-Gal4* driver. Scale bar: 20μm. Histograms: the horizontal bar indicates the mean, each dot represents an independent animal, except in panel K. Survival plots show a pool of at least 3 independent experiments. Statistics: ordinary one-way ANOVA followed by a Dunnett’s multiple comparisons test (B,C,E); Mann-Whitney test (F,K); Welch two sample t-test (G); CoxPH test followed by bonferonni corrections (H,I). ns, not significant; p ≥ 0.05, ^∗^ for *P* between 0.01 and 0.05; ^∗∗^ for *P* between 0.001 and 0.01, ^∗∗∗^ for *P* between 0.0001 and 0.001, ^∗∗∗∗^ for p ≤ 0.0001.

The main function of trachea is tissue oxygenation and CO_2_ disposal^29^. As expected, the reduced tracheation observed in *Tot* mutant flies resulted in lower tissue oxygenation, as measured by a transgene expressing a fluorescent oxygen sensor^30^ (**Fig. 3F**). Furthermore, flow-through respirometry revealed that *Tot^XMAZ^* flies consumed less O_2_ and produced less CO_2_ than wild-type (**Fig. 3G**). These results suggested that O_2_ delivery may limit the ability of *Tot*-deficient flies to cope with stress. To test this hypothesis, we exposed flies to osmotic stress at varying oxygen pressures. Hypoxia increased the susceptibility of both wild-type and *Tot^XMAZ^* flies to osmotic stress (**Fig. 3H**), highlighting the importance of oxygenation to resist this stress. Strikingly, hyperoxia partially rescued *Tot^XMAZ^* susceptibility to osmotic stress (**Fig. 3I**). These data demonstrate that Turandot-mediated support of oxygen delivery by trachea is critical to survive environmental stress.

Because *Tot*-deficient flies had reduced terminal tracheal cells, we hypothesized that these cells may undergo apoptosis. Consistent with this, we observed increased cleaved caspase-3 and TUNEL staining in trachea of *Tot*-deficient animals upon osmotic stress, as compared to wild-type flies (**Fig. 3J-L**). Conversely, expression of the apoptosis inhibitor p35^31^ in trachea restored wild-type TTC numbers in *Tot^AZX^* animals (**Fig. 3M**) and restored resilience to osmotic stress (**Fig. 3N**). These results show that Turandot proteins maintain oxygen supply upon stress by preventing tracheal cell apoptosis.

### Turandots prevent PS-dependent tracheal cell killing

Since TTC appeared to die from apoptosis in *Tot*-deficient flies, we looked at the presence of phosphatidylserine (PS), an early marker of cell death^13, 14^. PS is usually confined to the cytoplasmic leaflet of the membrane but becomes exposed upon apoptosis. PS asymmetry is maintained by flippases of the P4-ATPase family, while scramblases promote PS exposure on the outer layer of the membrane^32, 33^. A transgene expressing Annexin-V-GFP revealed increased levels of PS at the surface of tracheal membranes of *Tot*-deficient flies upon osmotic stress, as compared to wild-type flies (**Extended Data Fig. 4a,b**). Unexpectedly, we found that both wild-type and *Tot* mutant trachea spontaneously exposed measurable amounts of PS compared to other tissues, even without external challenge (**Fig. 4A and Extended Data Fig. 4c**). Strong Annexin-V staining was restricted to trachea and some muscles (**Extended Data Fig. 4 d,e**). We then explored whether the high PS exposure on tracheal cells could explain their susceptibility to stress. Overexpressing the mouse scramblase *xkr8* in trachea of wild-type flies to increase PS exposure reduced TTC numbers and resilience to osmotic stress (**Fig. 4B and 4C).** Conversely, lowering levels of PS exposed on trachea by knocking down *scramblase 1* in *Tot*-deficient flies restored normal TTC numbers and reduced susceptibility to stress (**Fig. 4D and 4E**). Collectively, these results indicate that high constitutive PS exposure on trachea contributes to their vulnerability to stressful conditions.

**Figure 4:**
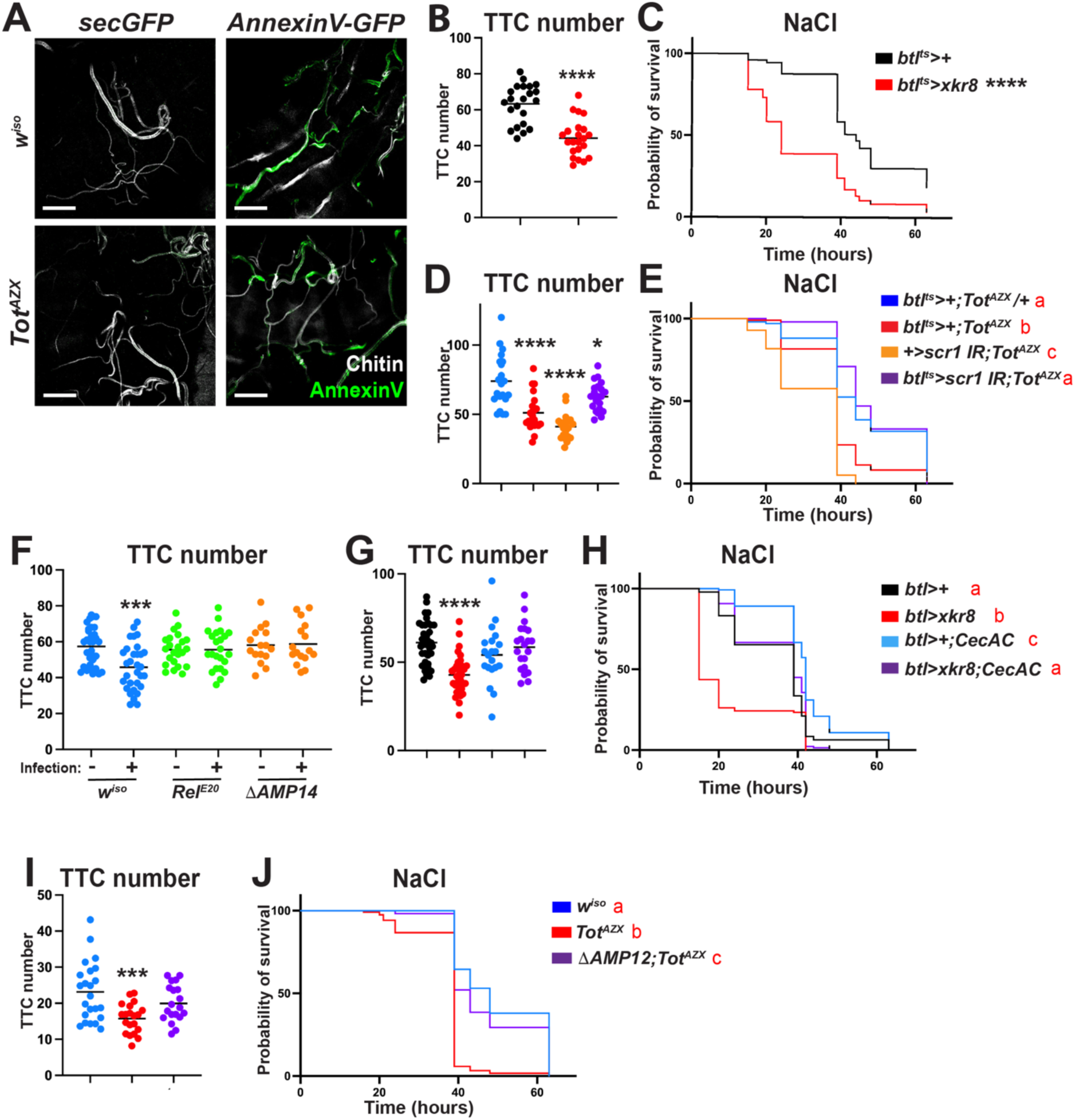
Turandots prevent PS-dependent tracheal cell killing by AMPs. **A**. Immunostaining of trachea from control (upper panels) and *Tot*-deficient flies (lower panels) overexpressing a secreted GFP (secGFP) (left) or Annexin-V-GFP (right) in the fat body. Blue, Chitin; Green, GFP; Scale bar: 20μm. **B,C**. Quantification of TTC numbers (**B**) and survival to osmotic stress (**C**) of wild-type flies overexpressing *xkr8* in trachea with the *btl^ts^* driver. **D,E**. Quantification of TTC numbers (**D**) and survival to osmotic stress (**E**) of *Tot^AZX^* flies over-expressing *scramblase 1* RNAi in trachea with the *btl^ts^* driver. (**F**) Quantification of TTC numbers of *w^iso^* (blue), *Rel^E^*^20^ (green) and *ΔAMP14* (orange) flies unchallenged or injured with heat-killed bacteria. **G,H**. A deletion removing four *Cecropin* genes (*CecA*C) rescues TTC numbers (**G**) and survival to osmotic stress of flies overexpressing *xkr8*. **I,J.** Quantification of TTC number (**I**) and survival to osmotic stress (**J**) of *w^iso^* (blue), *Tot^AZX^* (red) and *ΔAMP12;Tot^AZX^* (purple) flies. Scale bar: 20μm. Histograms: the horizontal bar indicates the mean, each dot represents an independent animal. Survival plots show a pool of at least 3 independent experiments. Statistics: ordinary one-way ANOVA followed by a Dunnett’s multiple comparisons test (B,F,G,I); Mann-Whitney test (D); CoxPH test followed by bonferonni correction (C,E,H,J). ns, not significant; p ≥ 0.05, ^∗^ for *P* between 0.01 and 0.05; ^∗∗∗^ for *P* between 0.0001 and 0.001, ^∗∗∗∗^ for p ≤ 0.0001.

### TotA prevents PS-dependent apoptosis of trachea by antimicrobial peptides

As PS is negatively charged, exposure on the outer leaflet is expected to sensitize cells to damage by cationic pore-forming AMPs^1, 11, 34^. We hypothesized that Tot may promote tracheal survival by preventing AMP-dependent cell death. To test this model, we monitored tracheal morphology in immune-challenged flies expressing high levels of AMPs. Infection with heat-killed bacteria decreased TTC number in wild-type animals, showing that the innate immune response adversely impacts trachea survival. Strikingly, neither Imd pathway deficient flies (*Relish^E^*^20^)^35^, that fail to express most AMPs, nor flies lacking 14 AMP genes *(τ.AMP14*)^36^, showed a reduction in TTC number after infection with heat-killed bacteria (**Fig. 4F**). These observations strongly suggested that AMPs directly kill tracheal cells during the immune response. We next tested whether tracheal killing by AMPs relied on PS exposure. The reduction of TTC numbers and the vulnerability to stress of flies displaying high PS exposure due to the overexpression of *xkr8* in trachea was rescued in absence of Cecropins, a major class of AMPs^36, 37^ **(Fig. 4G and 4H).** These data reveal that AMPs kill tracheal cells in a PS-dependent manner, leading to reduced resilience to osmotic stress. Assuming that Tots protect against AMP cytotoxic activity, we expected that removing AMP genes would be sufficient to reduce the vulnerability of *Tot* flies to stress. To test this notion, we generated a fly line simultaneously lacking six Tots and 12 AMPs and monitored tracheation and stress resilience in these flies. Strikingly, tracheation and resilience to osmotic stress in *τ.AMP12, Tot^AZX^* flies were similar to those observed in wild-type flies (**Fig. 4I and 4J**). We conclude that Tot proteins mitigate damage caused by AMPs to tracheal cells exposing PS.

### TotA prevents AMP-dependent pore formation

Previous studies have shown that AMPs can be cytotoxic to host cells in certain contexts^1, 3^.We therefore explored whether Tot proteins could protect host cells from AMP activity *in vitro*, focusing on *TotA*. Electrophysiology recordings on artificial lipid bilayers mimicking eukaryotic membranes confirmed that the antibacterial peptides Cecropin A (CecA) made transient pores in PS-rich membranes^38–40^ (**Fig. 5A**). Strikingly, addition of TotA abrogated CecA-induced pore formation (**Fig. 5B and 5C**). This result was confirmed in a liposome leakage assay (**Fig. 5D**). While CecA alone induced liposome permeabilization, addition of TotA was sufficient to abolish CecA-induced dye leakage. This protective role of TotA was not restricted to CecA. Indeed, TotA was able to prevent liposome leakage caused by Melittin, a potent pore-forming honey bee toxin with AMP activity^41^ (**Fig. 5E**).

**Figure 5:**
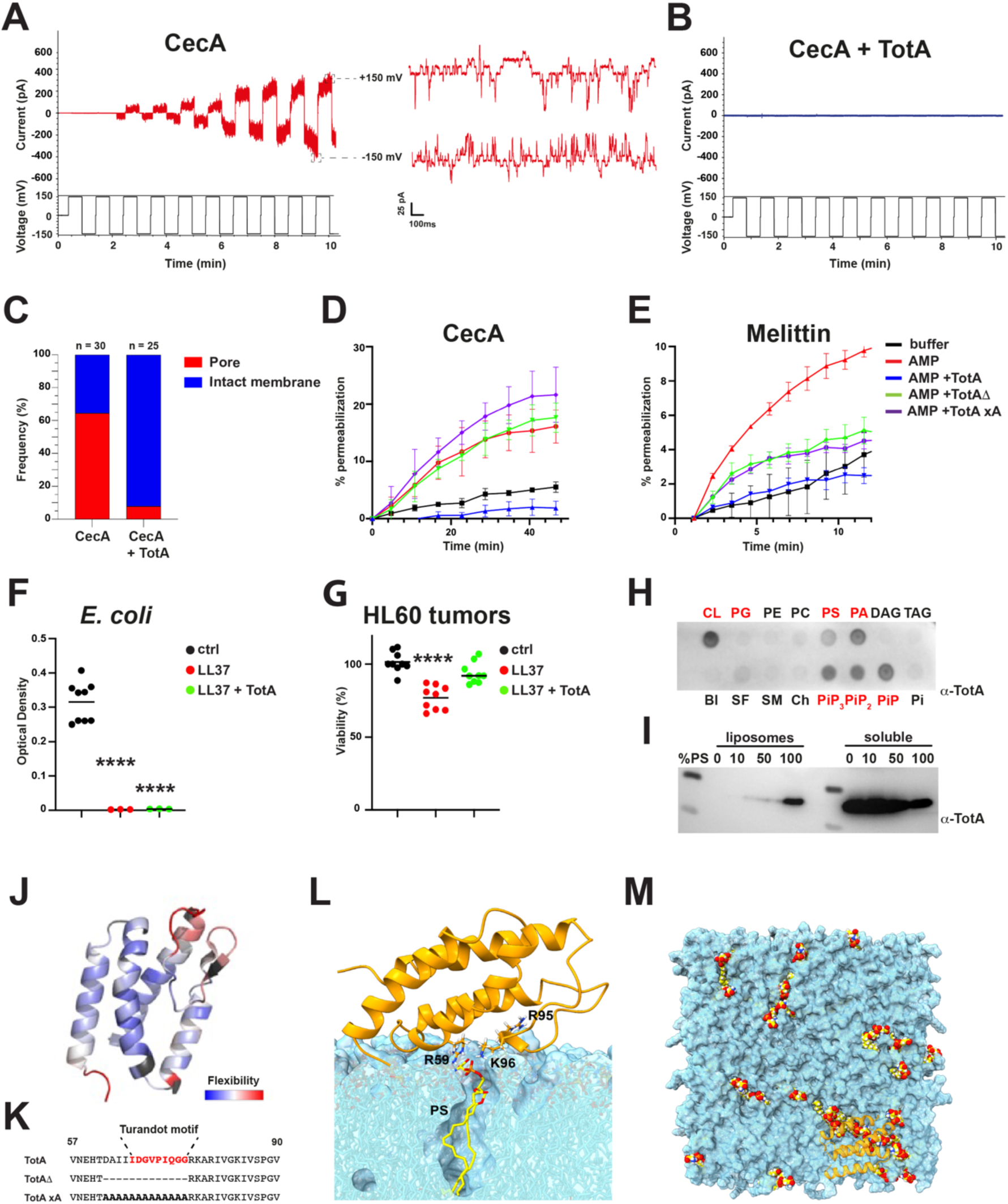
TotA prevents pore formation by AMPs. **(A-C)**. Current recordings through lipid bilayer after addition of CecA (**A**) or CecA + TotA (1:2) (**B**). Right panels in **A** show magnified current recordings at positive (upper panel) and negative (lower panel) voltages. (**C**). Quantification of pore formation (red) and intact membranes (blue) after addition of CecA (left) or CecA + TotA (right) to artificial bilayers in 3 independent experiments. (**D**-**E**) Kinetics of calcein leakage from liposomes incubated with buffer (black), CecA (**D**) or Melittin (**E**) and buffer (red), or 1μM TotA (blue), TotAΔ (green) or TotAxA (purple). Lines represent averaged values of 3 (cecA) or 2 (melittin) independent experiments with standard deviations. (**F,G**) Viability of *E. coli* (**F**) and HL60 cells (**G**) after incubation with buffer (white), LL37 (orange) or TotA + LL37 (blue). Shown are means (bar) and individual values of 3 independent experiments. (**H**) Representative membrane of a lipid overlay assay incubated with TotA and revealed by anti-TotA antibody. CL, cardiolipin; PG, phosphatidylglycerol ; PE, phosphatidylethanolamine; PC, phosphatidylcholine; PS, phosphatidylserine; PA, phosphatidic acid; DAG, diacylglycerol; TAG, triacylglycerol; PI, phosphatidylinositol; PiP, phosphatidylinositol (4)-phosphate; PiP_2_, phosphatidylinositol (4,5)-bisphosphate; PiP_3_ phosphatidylinositol (3,4,5)-trisphosphate; Ch, cholesterol; SM, sphingomyelin; SF, 3-sulfogalactosylceramide; Bl, blank. Negatively charged lipids are highlighted in red. (**I**) Anti-TotA Western blot analysis of liposome-containing and soluble fractions after incubation of TotA with liposomes containing increasing amounts of PS. (**J**) TotA structure as determined by NMR. Mapping of ^1^H-^15^N heteronuclear NOE onto the NMR structure of TotA, where red hints at more dynamic regions. (**K**) Sequences of TotA from position 57 to 90 containing the Turandot motif (in red) and two variants with this motif deleted (TotAΔ) or alanine-replaced (TotAxA). (**L**) Most representative binding conformation of TotA (orange cartoon) interacting with PS (yellow and red sticks) during the MD simulations. Most relevant charged residues for PS specificity are shown in sticks. (Blue, PE and PC). (**M**) Representative snapshot of the MD simulation showing a lipid bilayer (blue) with some PS lipids (yellow and red spheres) clustering beneath TotA (orange cartoon).

In addition to its antimicrobial activity, the human AMP LL37 has potent antitumoral properties, being capable of killing HL60 human leukemia cells^42^. We explored the impact of TotA on both LL37 microbicidal and antitumoral activities. TotA did not affect LL37 antibacterial activity on *Escherichia coli* (**Fig. 5F**). However, HL60 cells treated with a mixture of LL37 and TotA had higher viability than cells treated with LL37 only, showing that TotA inhibited LL37 activity specifically against eukaryotic cells (**Fig. 5G**). These data indicate that TotA directly inhibits the activity of a broad range of AMPs specifically on host cells.

### TotA protects host cells from AMPs by sequestering PS

To address the biophysical mechanism underlying AMP inhibition by TotA, we next tested whether TotA could sequester CecA or LL37 in solution. Isothermal titration calorimetry did not reveal any interaction between these AMPs and TotA in solution (**Extended Data Fig. 5a-d**). Therefore, we explored whether TotA could interact with membrane bilayers. Electrophysiology recordings revealed a transient membrane disruption after TotA addition, suggesting a dynamic interaction of TotA with membranes. (**Extended Data Fig. 5e).** We next tested whether TotA could bind to phospholipids using a lipid overlay assay. Strikingly, TotA binding was restricted to negatively charged phospholipids, with the exception of the bacterial lipid phosphatidylglycerol (**Fig. 5H**). Accordingly, TotA protein bound to liposomes in a PS-dependent manner, further confirming that TotA interacts with negatively-charged phospholipids (**Fig. 5I**).

We then took a structural approach to explore how TotA interacted with membranes. The solution structure of TotA as determined by NMR revealed a four-helix bundle with an internal disordered loop between helix 3 and 4 (**Fig. 5J**). This loop was less defined than the rest of the protein, likely because several residues at and around the loop were dynamic across a wide range of timescales. (**Fig. 5J and Extended Data Fig. 5f-h**). This flexible loop contains the so-called Turandot motif (*[I/V]-D-G-v-p-x-Q-G-G*), which is shared by all Tot members^4^ (**Fig. 5K**). We hypothesized that this flexible loop may be important for TotA-mediated AMP inhibition. To test this hypothesis, we designed two protein variants where the loop was deleted (TotAΔ) or entirely replaced with alanines (TotAxA) (**Fig. 5K**). Strikingly, both TotA variants were unable to prevent AMP-induced liposome lysis by CecA and Melittin, in line with the idea that the loop is necessary for AMP inhibition (**Fig. 5D and 5E**).

We then performed molecular dynamics (MD) simulations of TotA by placing it on the surface of a lipid bilayer model. TotA spontaneously reoriented itself on the membrane, with the Turandot motif loop facing the membrane surface (**Fig. 5L and Supplementary Information**). Strikingly, TotA displayed longer residence times around PS when compared to phosphatidylethanolamine (PE) or phosphatidylcholine (PC) lipids, despite PS representing only 10% of the membrane lipid composition (**Extended Data Figure 6a**). Three TotA residues, R59 in helix 2 and R95 and K96 adjacent to the Turandot motif, individually contributed the most to PS binding during the simulations (**Fig. 5L**), as assessed by their residence times and occupancy percentage (**Extended Data Figure 6a and b**). This set of interactions triggered a clustering of PS around TotA (**Fig. 5M and Supplementary Information**). In contrast TotA did not co-localize as frequently with PC and PE (**Extended Data Figure 6c and 6d)**. Collectively, these results suggest that TotA can sequester PS, thereby preventing their interaction with AMPs and subsequently preserving membrane integrity.

## Discussion

In this article, we provide evidence that Turandot proteins promote stress resilience in *Drosophila* by protecting host tissues from AMP-dependent lysis. Strikingly, we observed that TotA neutralizes the pore forming activity of CecA and LL37 on host cells without impacting their microbicidal activity. TotA binds *in vivo* to tracheal cells and can interact with PS-enriched artificial membranes *in vitro*. MD simulations and lipid binding assays show that TotA preferentially interacts with PS, clustering and masking this phospholipid. We hypothesize that PS recruitment and sequestration by TotA shields PS from AMPs, therefore preventing AMP recruitment at the membrane. This would explain the broad protective effect of TotA against phylogenetically distant AMPs. By selectively inhibiting pore formation on eukaryotic cells, Turandot proteins increase AMP selectivity, allowing production of microbicidal AMPs at high concentrations while reducing collateral damage to host tissues.

Surprisingly, Annexin-V staining reveals that tracheal cells constitutively expose high levels of PS in the outer leaflet of the membrane. High PS exposure reduces the asymmetry between the inner and outer leaflet of the membrane and is thought to facilitate cell deformation and prevent shear stress^43, 44^. This could be especially important in tracheal branches that are intimately attached to motile tissues, including the muscles and gut, and are constantly exposed to mechanical stress^29^. We believe that Tots emerged through evolution to protect trachea and other tissues that are sensitized to lysis by AMPs due to reduced membrane asymmetry.

Restoration of homeostasis upon stress is energetically costly and requires increased respiration. That lethality of *Tot*-deficient flies with reduced tracheation can be rescued by hyperoxia illustrates that tissue oxygenation by trachea is critical to survive stress. We suspect that both basal and stress-induced Tot expression by epithelia and the fat body protect the respiratory epithelium. During pupariation, larval trachea undergoes histolysis and adult trachea arise from pupal progenitors. Interestingly, both Tots and AMPs are highly expressed during this stage^4, 45^. AMPs are expected to play a prophylactic role during this stage to prevent infection by bacteria escaping the gut during metamorphosis^46^. High Tot expression during metamorphosis would in turn protect trachea from AMPs at this critical stage.

Reports increasingly indicate that AMPs and other cationic peptides can be cytotoxic to host cells in certain contexts^10–12^, notably in neurodegeneration^47^. High PS exposure has also been associated with axon degeneration^33^, Here we show that exposure of PS^48, 49^ upon stress or in certain cell types sensitizes them to cationic AMPs. It is therefore tempting to speculate that high PS exposure in neurons could sensitize this tissue to AMP-mediated killing. To our knowledge, our study is the first to identify a class of molecules protecting animal cells from the action of AMPs. Similar mechanisms might exist in other organisms, as suggested by an *in vitro* study showing that two well-known antimicrobial peptides, LL37 and HNP1b, cooperate to kill bacteria more efficiently while minimizing mammalian cell membrane lysis^50^. Identification of factors such as Turandots that protect host cells from AMPs is therefore of therapeutic interest in several contexts, including neurodegeneration.

## Material and methods

### Drosophila stocks and genetics

Flies were raised on Yeast-Cornmeal food (6% cornmeal, 6% yeast, 0.62% agar, 0.1% fruit juice, that was supplemented with 10.6g/L Moldex and 4.9ml/L propionic acid) at 25°C. Experiments were performed on 5-10 days old animals at 25°C, unless otherwise stated. Animals were bred and maintained at a low population density in vials and flipped twice a week. Isogenic *w*^1118^ Drosdel flies^51^ (*w^iso^*) were used as wild-type. *TotM* and *TotX* mutants were generated as previously described^52^. Briefly, *w^iso^* embryos were injected with a mixture of recombinant Cas9 (Invitrogen) and a gRNA targeting the *TotM* coding sequence (ACTTATCGTAGAAAGTGACCAGG) or the *TotX* coding sequence (GTTCAAGTTATGAGGAACACAGG), respectively. *TotAZ^sk^*^6^ line was created as previously described in^53^. This mutation was subsequently backcrossed in the *w^iso^* background. To generate *Tot^AZX^* stock, *TotAZ^sk^*^6^ embryos were injected with a mixture of recombinant Cas9 (Invitrogen) and a gRNA targeting TotX coding sequence (GTTCAAGTTATGAGGAACACAGG). *Tot^AZX^*mutations were then combined with *TotM* mutation (on the second chromosome) to create the *Tot^XMAZ^* line. All *Tot^AZX^* mutations were located on the third chromosome and this mutant almost phenocopied *Tot^XMAZ^*. Additionally, most of the transgenes and genomic deletions used in our genetic analysis were located on the second chromosome. We therefore decided to use *Tot^AZX^* instead of *Tot^XMAZ^* in this study in order to simplify the genetic schemes. U*AS-TotA-HA* and *UAS-HA-TotA* lines were generated by phiC3-mediated recombineering. Stocks used in this study are listed in **Extended Data Table 1**.

### Cloning and DNA constructs

Cloning was performed by Gibson assembly (New England Biolabs), following the manufacturer’s instructions. *TotA* coding sequence was cloned into the pUASt-attb-GFP vector and a HA tag was added in C-term. A sequence containing the TotA 5’UTR, TotA signal peptide, 3 times a HA-tag and TotA CDS was ordered from Genewiz and subsequently cloned into the pUASt-attb-GFP vector to create the UAS-HA-TotA construct. For recombinant protein expression, a codon-optimized version of TotA fused to a Tobacco Etch Virus (TEV) protease cleavage site was ordered from Twist Bioscience and cloned into the pET29b vector. TotAxA and TotAΔ were made by GenScript, using pET29b-TotA as a template.

### Recombinant protein and antibody production

TotA, TotAxA and TotAΔ were expressed overnight at 18°C in Rosetta2 *E. coli* (DE3, Novagen). Cells were lysed by sonication in 700 mM NaCl;20 mM HEPES 7.5 containing 1 protease inhibitor tablet (Roche) + 5ul Turbonuclease. TotA was purified using HisPur Ni-NTA Resin beads (ThermoFisher). MBP protein was cut using super TEV protease and both proteins were removed using HisPur Ni-NTA Resin beads. TotA was further purified by incubation with Amylose Resin (New England Biolabs) followed by size-exclusion chromatography on a HiLoad superdex75 16/60 column (GE healthcare) and concentrated in 150 mM NaCl; 20 mM HEPES 7.5. TotA was used as immunogen to produce rabbit anti-TotA antibody (made by GenScript).

### Microbial cultures

Bacteria were cultured overnight on a shaking plate at 180 RPM. The following morning, they were pelleted by centrifugation (4000 RPM at 4°C) and the bacterial pellets were diluted to the desired optical density at 600 nm (OD_600_). *Pectobacterium carotovorum carotovorum 15* (*Ecc15*) and *Micrococcus luteus* were grown in LB medium at 29°C. *Enterococcus faecalis* was grown in GHI medium at 37°C. For experiments with heat killed bacteria, bacterial pellets were cyclically (4 times) heated at 95°C and frozen at -20°C. *Drosophila C Virus* (*DCV*) stock was kindly provided by Prof. Carla Saleh.

### Survival experiments

All experiments were done on 5-to 7-days old adult female flies. Systemic infections with *Ecc15*, *M. luteus* and *E. faecalis* were performed as follow^54^: flies were pricked in the thorax with a needle previously dipped into a concentrated bacterial pellet at OD_600_:200 (*Ecc15* and *M. luteus*) or OD_600_:5 (*E. faecalis*). Infected flies were maintained at 25°C (*E. faecalis*) or 29°C (*Ecc15* and *M. luteus*) and survival was recorded daily. Flies were flipped into fresh vials every 2-3 days. Systemic infections with DCV were performed by injecting 50 nL of 2*10^5^ TCID_50_/mL into the thorax of female adult flies using a nanoinjector (Drummond) and glass capillary needles.

Survivals to abiotic stresses were performed as followed: osmotic stress experiments were performed by feeding fly food supplemented with NaCl (final concentration: 4%). Flies carrying Gal4 or Gal4^ts^ overexpressing transgenes were raised at 25°C, kept for 3days at 25°C, transferred for 2-4days at 29°C and exposed to an osmotic stress at 29°C. Starvation experiments were performed by keeping flies on 1.8% agar. For heat stress experiments, flies were kept in a 34°C incubator. For hypoxic/hyperoxic experiments, flies were maintained in an incubator with controlled oxygen levels (5% O_2_ for hypoxia of 60% O_2_ for hyperoxia).

### Western Blot

Total hemolymph was harvested by bleeding ten L3 wandering larvae in 100 μL of Phosphate Buffer Saline (PBS) supplemented with Complete Inhibitor (Roche, diluted 1/50), PMSF (1 mM) and PTU (1 μM). A step of centrifugation (5 minutes at 500 x g) was done to discard hemocytes. 10 μg of proteins was denatured (2 minutes at 80°C) and then separated on a Novex 10-20% precast Tricine Gel, and transferred to a nitrocellulose membrane (Invitrogen iBlot). The membrane was blocked in 5% non-fat dry milk in PBS containing 0.1% Tween-20 for 1 hour and then incubated overnight at 4°C with a rabbit anti-TotA (1:1000 dilution) antibody. Goat anti-rabbit-HRP secondary antibody (Jackson ImmunoResearch) diluted at 1:15000 was incubated for 1h at room temperature. Bound antibodies were detected using SURESIGNAL Western Substrate (lubioScience). The membrane was imaged on a ChemiDoc XRS+ (BioRad).

### Immunostaining and trachea imaging

Guts were dissected in PBS and fixed in 8% paraformaldehyde for 15 minutes at room temperature. The tissues were subsequently rinsed two times with PBS containing 0.1% Triton X-100, blocked for 1 hour in PBS containing 1% of goat serum and 2% BSA, and incubated at 4°C in the blocking solution containing an anti-GFP (2hours, 1:3000), anti-HA (overnight, 1:500) or anti-cleaved Caspase3 primary antibody (overnight, 1:500). The tissues were subsequently washed and incubated with a secondary antibody (1:4000) for 1 hour at room temperature. After extensive washes, tissues were mounted in glycerol. Anti-TotA staining was done following a similar protocol without adding Triton X-100. To image AnnevinV-GFP, guts were dissected in AnnexinV buffer (Hepes 10mM, NaCl 150mM, KCl 5mM, MgCl_2_ 5mM, CaCl_2_ 1.8 mM) and subsequently stained following the abovementioned protocol. Anti-TotA staining of fat bodies was performed on dissected carcasses. Tissues were fixed for 45 min, permeabilized in PBS-0.1% Triton X-100 for 1 hour, blocked for 1 hour in PBS containing 1% of goat serum and 2% BSA, and incubated overnight at 4°C in the blocking solution containing anti-TotA antibody (1:3000). The tissues were subsequently washed and incubated with a secondary antibody (1:4000) for 1 hour at room temperature. After extensive washes, tissues were mounted in glycerol. Tracheation of guts was assessed by imaging the autofluorescent chitin lining the tracheal lumen. The dityrosine bonds chitin fluoresce under UV excitation, allowing chitin to be imaged using a DAPI filter set. Flies were starved for two hours before guts were dissected and fixed in 8% paraformaldehyde for 45 minutes at room temperature. Guts were subsequently washed 3 times 15min in PBS and mounted in glycerol. Guts were imaged on a LSM700 microscope (Zeiss) using the DAPI channel.

### Tunel Staining

Guts from adult female flies were dissected in PBS and fixed in 4% paraformaldehyde for 15 minutes at room temperature. The guts were subsequently rinsed three times with PBS and permeabilized for 2 minutes in PBS+0.1% Triton X-100 + 0.1% Sodium citrate tribasic dihydrate, and rinsed three times in PBS. Guts were then incubated for 1 hour with reagents from the *In Situ Cell Death Detection Kit, TMR red* (Roche). Tissues were washed three times and mounted in Dako Fluorescence Mounting Medium (Agilent).

### Tissue oxygenation determination

Oxygenation of Drosophila tissues was determined using a genetically-encoded fluorescent probe (nls-timer) as described^30^. Guts and Malpighian tubules were dissected and fixed in 4% paraformaldehyde for 45 minutes and mounted in glycerol. Images were acquired in the GFP and RFP channels on a Leica M205 FA fluorescent stereomicroscope. Ratiometric analysis of red and green nuclei fluorescence was performed using ImageJ to determine tissue oxygenation.

### Image analysis and TTC quantification

Image quantifications were performed using ImageJ software. Trachea coverage was determined by measuring the surface of trachea divided by the area of the R2 gut region. Terminal tracheal cells cellular bodies, as defined by the characteristic shape of the chitinous lumen, were manually counted in the R2 region of the midgut.

### Respirometry

Respiration in flies was measured using a stop-flow gas-exchange system (Q-Box RP1LP Low Range Respiration, Qubit Systems, Ontario, Canada, K7M 3L5). Eight female flies were put into an airtight glass tube and supplied with food. Each tube was provided with CO_2_-free air while ‘spent’ air was simultaneously flushed through the system and analyzed for its CO_2_ and O_2_ content. In this way, evolved CO_2_ and consumed O_2_ were measured for each tube every ∼ 44 minutes for a duration of 12h-16h.

### NMR spectroscopy

Recombinant proteins for NMR spectroscopy were prepared with the isotope enrichment scheme appropriate for each experiment in MES buffer pH 6 with 100 mM NaCl and 10% ^2^H_2_O. Protein concentrations were around 300 µM for NMR titrations and close to 1 mM for experiments aimed at resonance assignment, structure determination, and ^15^N relaxation analysis. All NMR experiments were carried out in a 18.8 T (800 MHz ^1^H Larmor frequency) Bruker spectrometer equipped with a CPTC ^1^H,^13^C,^15^N 5 mm cryoprobe and an Avance Neo console. Backbone (H, N, CA, and C) and CB resonances were assigned using a standard procedure based in conventional 3D HNCA, HN(CO)CA, HNCO, HN(CA)CO, CBCA(CO)NH and HNCACB spectra, further assisted by ^15^N-resolved TOCSY and NOESY^55^. Experiments for sidechain assignment and structure calculation entailed ^15^N-resolved NOESY, ^15^N-resolved TOCSY, HCCH-TOCSY, ^13^C-resolved NOESY, HNHA, 2D TOCSY and 2D NOESY spectra. All spectra were acquired and processed using Bruker’s Topsin 4.0 software. Backbone and sidechain assignments were obtained through manual spectral analysis assisted by the program CARA^56^. Analysis of NOESY spectra and structure calculations were performed semiautomatically using UNIO’s ATNOS/CANDID module coupled to Cyana^57, 58^ using NOEs and dihedral angles derived from chemical shifts with Talos-n^59^. NMR structure statistics are provided in **Extended Data Table 2**. The solved protein structure was deposited in the PDB under ID 8PBV and NMR chemical shifts at the BMRB under ID 34825.

For studies of ^15^N relaxation, heteronuclear ^1^H-^15^N NOEs were measured using an interleaved, phase-sensitive gradient-enhanced version; and ^15^N T_1_ and T_2_ were measured via conventional pseudo-3D experiments that apply respectively inversion recovery or CPMG sequences onto an HSQC spectrum, both using gradient selection, water suppression, and decoupling during acquisition. All spectra for measurement of ^15^N relaxation were acquired with 256 increments in the indirect dimension and 3 seconds of relaxation delay. For T_1_ measurements we used delays of 20, 50, 100, 200, 300, 400, 500, 700, 1000, 1200, 1500, 2000, 3000 and 4000 ms. For T_2_ measurements we used delays of 17, 34, 51, 68, 85, 102, 119, 136, 170, 204, 238, 272 and 340 ms. Relaxation rates were fitted from exponential decay models using the ad hoc module from the Sparky-NMRfam program^60^ on the peaks derived from our backbone assignment.

### Peptides

Purified CecA and Melittin (Sigma) were used in leakage experiments. Synthetic CecA and LL37 were purchased from GenicBio (purity> 95%).

### Isothermal Titration calorimetry

ITC experiments were performed on a Microcal PEAQ-ITC (Malvern Instruments). 13 injections of 2uL of a 10mM HEPES pH7.5 150mM NaCl solution of 2mM CecA or LL37 were injected 13 times in the chamber containing 200 μM TotA and heats of reaction were recorded. Control experiments were run where peptides were titrated in the buffer solution alone.

### Liposome preparation, binding and leakage experiments

Lipids (3 mg) were resuspended in chloroform and mixed at a 3:1:0.4 ratio (DOPE:DOPC:DOPS) with the exception of experiments involving melittin, in which DOPS was omitted, as this lipid inhibits the membranolytic activity of melittin. Chloroform was evaporated under a nitrogen stream and the lipid film was hydrated with 600 μL of 50 mM Tris pH=8, 100 mM NaCl containing 70 mM Calcein. After 5 freeze-thaw cycles, liposomes were extruded 20 times through a 0.2 μm pore-size filter. To remove un-entrapped calcein, liposomes were passed through two HiTrap columns at low flow rate. Fractions containing calcein-loaded liposomes were pooled, diluted 4 times and used for subsequent assays. For calcein leakage experiments, 10 μl liposomes were incubated in 90 μl 50 mM Tris pH=8, 100 mM NaCl containing 0.5 μM TotA, TotAxA or TotAϕλ, together with CecropinA and Melittin (Sigma-Aldrich, at a concentration of 1 μM and 8.5 nM, respectively) and fluorescence was recorded on an Infinite M Nano fluorospectrophotometer (Tecan) for 1 hour. Maximum release of Calcein was determined by addition of 0.1% triton-X100 and data were normalized as percentage of full release. For binding assays, 3 μM TotA was incubated with180 μL 1mg/mL liposomes containing DOPC and increasing amounts of DOPS for 30 minutes at room temperature and centrifuged at 100000 x g in a discontinuous sucrose gradient (35%:20%:10%) for 1 hour. Liposome fraction and soluble fractions were subjected to anti-TotA immunoblotting.

### Lipid overlay assay

Membrane lipid strips (Echelon Bioscience) were incubated with 0.5μg/mL TotA in PBS-0.1% Tween, 3%BSA for 1 hour at room temperature. TotA was then probed with an anti-totA antibody as described above.

### Single-channel bilayer experiments

1,2-diphytanoyl-*sn*-glycero-3-phosphocholine (DPhPC) and 1,2-diphytanoyl-*sn*-glycero-3-phospho-L-serine powder (DPhPS) (Avanti Polar Lipids Inc.,) were dissolved in octane (Sigma-Aldrich) to a final concentration of 8 mg/mL and either used pure or in mixture as specified in the figure caption. Single-channel recording experiments were performed on an Orbit 16 TC instrument (Nanion). Phospholipid membranes were formed across a MECA 16 recording chip that contains 16 circular microcavities (100 µm diameter) in a highly inert polymer. Each cavity contains an individual integrated Ag/AgCl microelectrode and can record 16 artificial lipid bilayers in parallel. The buffer (10 mM HEPES, 200 mM KCl, pH=7.4), the concentration of CecA and TotA was 8 µM and the temperature was set to 25°C for all experiments. Membranes were formed and their capacitance were recorded (**Extended Data Figure 5i**). The traces were recorded in Elements Data Reader (Elements) and further analyzed by Clampfit (Molecular device). Data was collected at 20 kHz sampling rate with a 10 kHz low-pass filter. Results, fitting, and graphs were produced in Prism (GraphPad) and figures were generated in Adobe Illustrator 2022 (Adobe).

### Killing Assays

A culture of *E. coli* grown overnight was diluted to OD600=0.001 and allowed to regrow for 2hrs at 29°C. 1 μl of this culture was diluted in 100μL LB containing 2μM LL37 in the absence or presence of 2.5 μM TotA. The plate was incubated at 25°C under intermittent agitation and OD was recorded every 10 minutes. For eukaryotic cell killing assay, 10000 HL60 cells were seeded in 100μL of RPMI supplemented with heat-inactivated FBS, HEPES and antibiotics in half-area 96-well plates (corning). The next day, 50 μL of supernatant was removed and LL37 and TotA were added at 40 and 80 μM final concentration, respectively. After 1h incubation at 37°C, 5 μL of MTT (Biotium) was dispensed in each well and further incubated at 37°C for 1h. 100 μL DMSO were then added and thoroughly mixed to dissolve crystals. Absorbance was read at 570nm and 630nm on an Infinite M nano spectrofluorometer (Tecan).

### Molecular Dynamics simulation

We obtained the structure of TotA from the AlphaFold Protein Structure Database (AFDB) ^61, 62^ under the UniProt accession code Q8IN44 ^63^. The models available in AFDB are computed based on the entire sequence of the target proteins, so it was necessary to remove the sequence peptide of TotA (residues 1 to 21) from the structure prior to preparing the simulation setup.

We processed the TotA model in the CHARMM-GUI web server^64^ to generate the input files for the MD simulation, using the CHARMM-GUI Membrane Builder tool^65^. We used the Positioning of Proteins in Membrane (PPM) tool from Orientations of Proteins and Membranes (OPM) database to orient TotA with respect to the membrane^66^. We generated a tetragonal box with TotA and a symmetric lipid bilayer with 10% DOPS, 30% DOPC, and 60% DOPE, resulting in a membrane with an area of 131 × 131 Å2. We solvated the system with TIP3P water^67^ and neutralized its liquid charges with counterions corresponding to a 0.15 M NaCl solution. Lastly, we generated all the necessary files to run the MD simulations with the CHARMM36m forcefield^68^.

We used GROMACS^69^ version 2022.1 to run and analyze the MD simulations. In all stages of the procedure, the cut-off radius for short-range electrostatic and van der Waals interactions was 12 Å; we used the particle mesh Ewald algorithm (PME)^70^ to calculate long-range electrostatic interactions. We restrained the length of covalent bonds involving hydrogen atoms using the LINCS algorithm^71^. Periodic boundary conditions were applied in all directions.

In the first stage of the simulation protocol, we minimized the potential energy of the system using the steepest descent algorithm, employing a 1000 kJ mol-1 nm-1 maximum force constant on the atoms as a convergence criterion. After minimization, we subjected the system to thermal equilibration in the canonical ensemble (NVT)^72^, to accommodate water and counterions around the protein and membrane. Initial atomic velocities were determined following a Maxwell-Boltzmann distribution corresponding to a temperature of 300 K and we equilibrated the system under these conditions for 0.25 ns with a 1 fs time step using the Berendsen thermostat^73^. During the equilibration stage, we restrained heavy-atom positions using a harmonic potential, with force constants of 4000 (backbone) and 2000 (sidechain) kJ mol-1 nm-2, respectively. The phosphorus atoms of the lipid molecules were also restrained on the Z-axis (direction of the normal vector at the membrane surface), as well as the dihedral angles of the double bonds in the fatty acid chains, with force constants of 1000 kJ mol-1 nm -2 and 1000 kJ mol-1 rad-2, respectively.

After controlling the temperature, we stabilized pressure and density in the isothermal-isobaric ensemble (NPT)^74^. The pressure was maintained at 1 bar using the Berendsen barostat^73^ with semi-isotropic pressure coupling and a 5 ps time constant. The systems were equilibrated for 1.625 ns, with position restraints released gradually.

The production dynamics were also performed in the NPT ensemble, changing only the thermostat used for Nosé-Hoover^75, 76^ and the barostat to Parrinello-Rahman^77, 78^. This stage lasted 1 μs, with a time step of 2 fs. All simulations had five replicas and only the last 500 ns of each simulation were used for the analyses.

We performed all lipid interaction analyses with the Python package PyLipID^79^. The distance cutoff for lipid interactions was set to 0.3 nm, and only residence times calculated with an estimated R2 > 0.97 were reported.

We used the Visual Molecular Dynamics (VMD)^80^ software for visual inspection of the simulations and recording of the movies. All protein and membrane images were generated with ChimeraX^81^. Density and violin plots were generated with the R programming language in RStudio^80, 81^.

### Statistical analysis

Each experiment was repeated independently at least three times. Survival curves included experiments with at least one cohort of 20 flies per condition. Survival analyses were performed using a Cox proportional hazards (CoxPH) model, with Bonferonni corrections for *P*-values when multiple comparisons were done. In this case, Statistics were represented using a compact letter display (CLD) graphical method: groups were assigned the same letter if they were not significantly different (p > 0.05). Quantification data were analyzed using a Mann-Whitney test, Welch two sample t-test, student t-test or ordinary one-way ANOVA with Dunnett’s multiple comparisons test, as stated in the figure legends. Error bars represent the standard deviation of the mean of replicate experiments (SD). *p*-values are represented in the figures by the following symbols: ns for p ≥ 0.05, ^∗^ for *P* between 0.01 and 0.05; ^∗∗^ for *P* between 0.001 and 0.01, ^∗∗∗^ for *P* between 0.0001 and 0.001, ^∗∗∗∗^ for p ≤ 0.0001. Survival statistics were represented using a compact letter display graphical technique: groups were assigned the same letter if they were not significantly different (P > 0.05). All survival statistics are summarized in **Extended Data Table 3**.

## Acknowledgements

We thank the Bloomington Stock Center in the USA and the Vienna Drosophila Resource Center for fly stocks; Jordi Casanova, Stefano de Renzis and Chun Han for providing fly strains; the BioImaging & Optics Platform (BIOP) in EPFL for confocal microscopy; the Protein Production & Structure Core Facility (PTPSP) for TotA production and NMR analysis; Hannah Westlake and Florent Masson for comments and editing on the manuscript. This work was supported by Sinergia SNSF grant CRSII5_186397, the SNSF project grant 310030_215073, PR00P3_193090 205321_192371 to Matteo Dal Peraro and Novartis Foundation 532114 awarded to Bruno Lemaitre, 532589 to Chan Cao.

## Authors Contributions

SR and BL designed the study and wrote the manuscript with inputs from all the authors. SR, AC, JBJ, LAA, CV, VR designed and performed experiments and analyzed data. JPB performed experiments and generated mutant and transgenic stocks. LAA performed the NMR experiments, solved TotA structure and wrote the manuscript. FM ran and analyzed the MD simulations. CV generated and analyzed respirometry data. SK provided mutants. MSD and MDP acquired funding and provide feedbacks. CC acquired funding and supervised experiments. BL acquired funding and supervised the project.

## Competing Interests Statement

The authors have no competing interests to declare.

Supplementary Information is available for this paper.

Correspondence and requests for materials should be addressed to Samuel Rommelaere. Reprints and permissions information is available at https://doi.org/10.6084/m9.figshare.23735694.

