## Supplemental Information for "A humoral stress response protects *Drosophila* tissues from antimicrobial peptides"

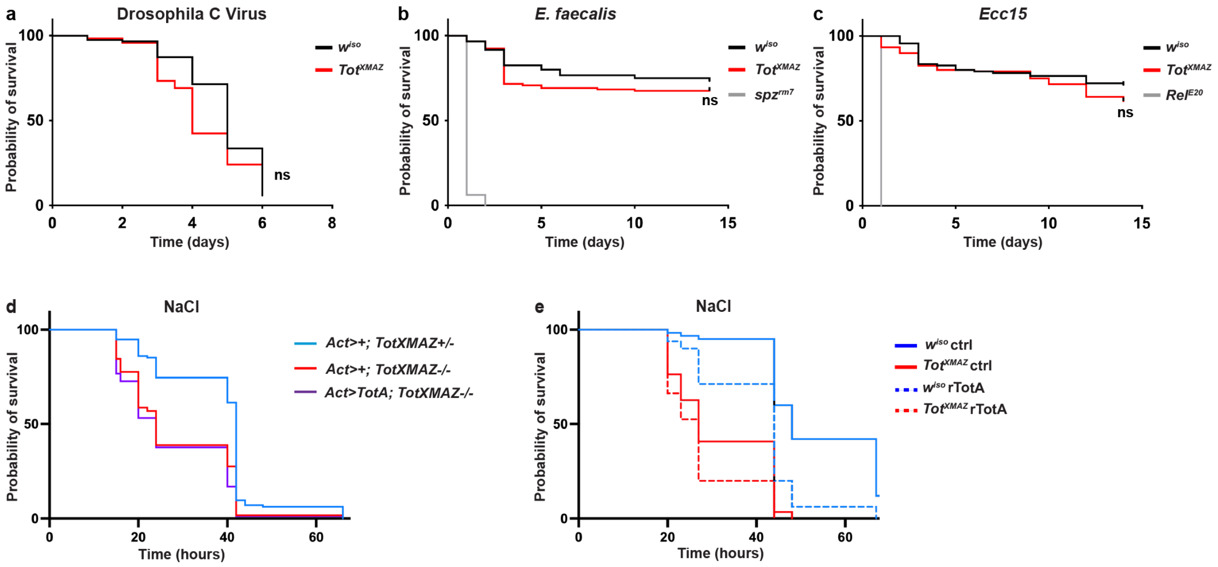

**Extended Data Figure 1**: **Additional survival phenotypes.** Turandot mutants are not strongly susceptible to microbial infections. *$w^{iso}$* (black) or *$Tot^{XMAZ}$* (red) female flies were systemically infected with Drosophila C virus (**a**), *Enterococcus faecalis* (**b**) or *Pectobacterium carotovora* (*Ecc15*) (c). *$Relish^{E20}$* or *$spz^{rm7}$* immune-deficient flies (grey) were used as positive controls in (**b**,**c**). ns, not significant (Cox-PH test). (**c**) Excessive amounts of TotA reduce resilence to stress. Survival to osmotic stress of *act>+, $Tot^{XMAZ}$/+* (red), *act>+, $Tot^{XMAZ}$* (blue) or upon ubiquitous overexpression of TotA in *$Tot^{XMAZ}$* background (*act>totA, $Tot^{XMAZ}$* purple). (**d**) Survival to osmotic stress of *$w^{iso}$* (blue) or *$Tot^{XMAZ}$* (red) flies upon injection of PBS (solid lines) or recombinant TotA (dashed lines).

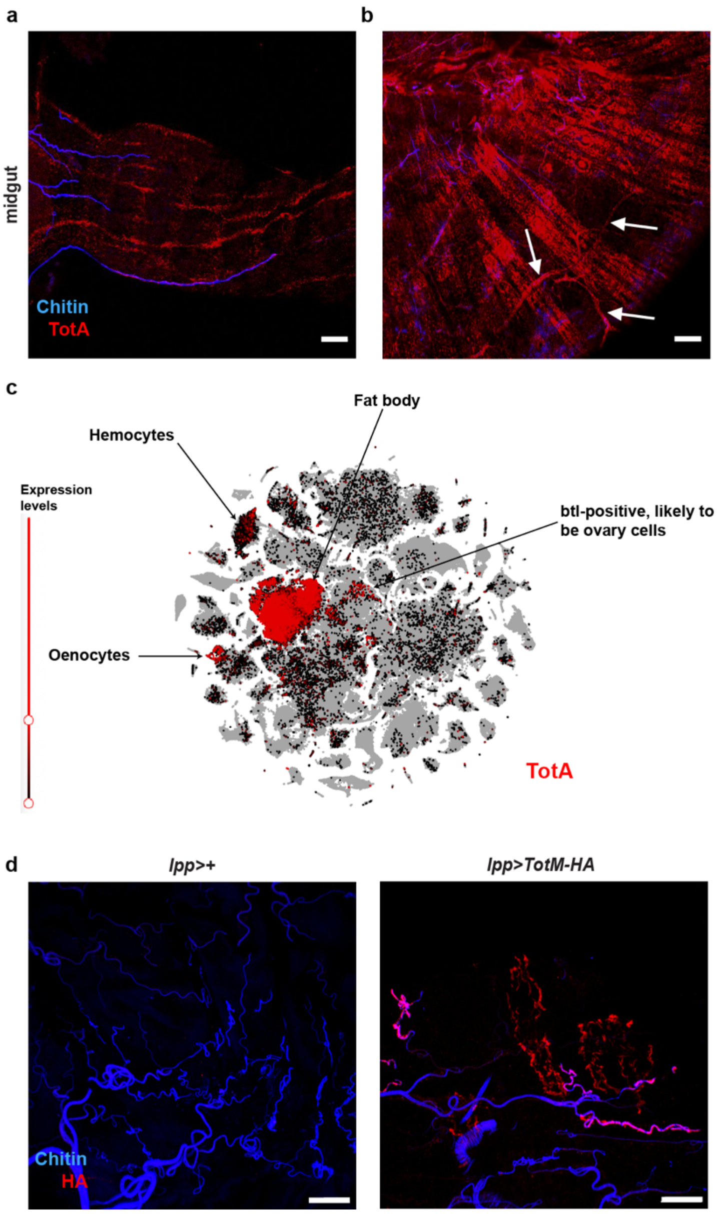

**Extended Data Figure 2**: **Fat-body-secreted Turandots bind to tracheas and visceral muscles.** (**a**,**b**) TotA localizes to the visceral muscles. Anti-TotA immunostainings of adult $w^{iso}$ midguts reveal localization to the longitudinal (**a**) and circular (**b**) visceral muscles as well as tracheas (arrows). Blue, chitin; Red, TotA. Scale bar: 20 μm. (**c**) The TotA gene is mostly expressed in the fat body. *t*-distributed stochastic neighbor embedding (t-SNE) plot representing single-cell expression of TotA (red). Shown are TotA expression levels as measured in the 10X Stringent Dataset extracted from FlyCellAtlas ([67]). TotA is mostly expressed in the fat body, in a subset of oenocytes and in hemocytes. Note that some btl-positive cells, which are likely to be ovary cells, express detectable levels of TotA. (**d**) Anti-HA immunostainings of adult *lpp*>+ (left) and *lpp*>*TotM-HA* (right) midguts. Blue, chitin; Red, HA. Scale bar: 20 μm.

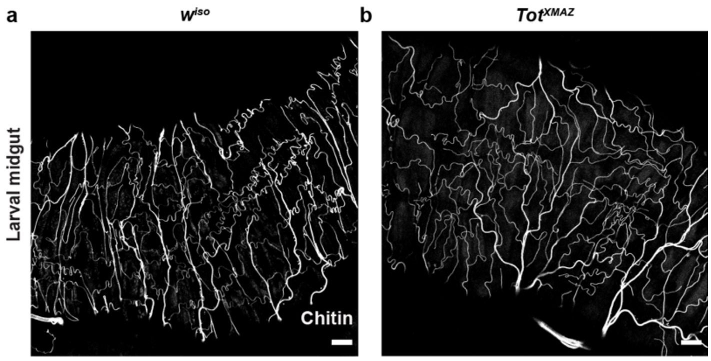

**Extended Data Figure 3: Larval gut trachea of *w*$^{iso}$ and *Tot*$^{XMAZ}$ larvae have comparable morphologies.** Representative pictures of larval gut trachea. Larval tracheas were imaged using chitin autofluorescence (white) of gut trachea of *w*$^{iso}$ (left) or *Tot*$^{XMAZ}$ (right) L3 larvae. Scale bar: 20 μm

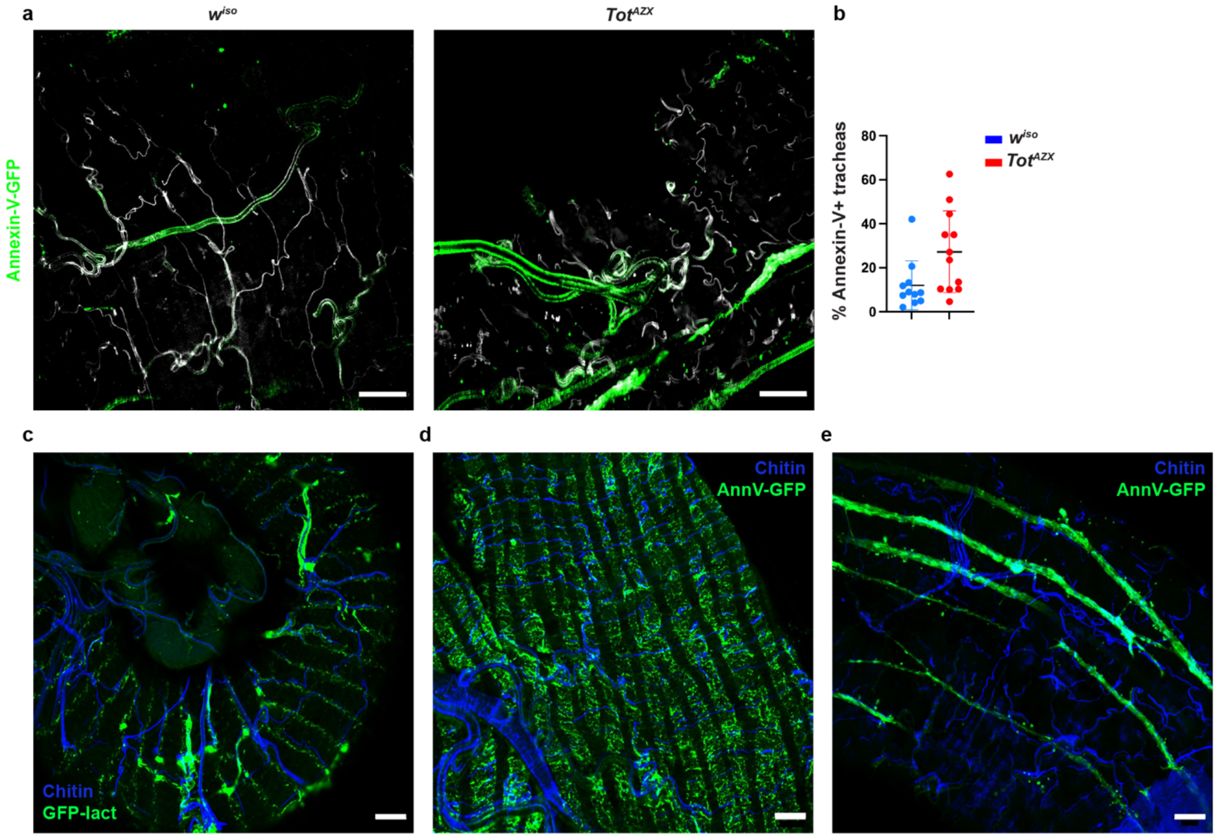

**Extended Data Figure 4: Labeling of extracellular phosphatidylserine in adult flies. (a).** Anti-GFP (green) immunostaining of trachea from control (left panel, *lpp>AnnexinV-GFP*) or Tot-deficient (right panel, *lpp>AnnexinV-GFP;Tot*$^{AZX}$) flies exposed to osmotic stress. (**b**) Quantification of the tracheal surface stained by AnnexinV in control (blue) and Tot deficient (red) flies fed NaCl food. (**c,d**)**.** Anti-GFP (green) immunostaining of trachea (**c**) and visceral muscles (**d**) from flies overexpressing the PS-binding protein lactadherin (*lpp>Lact-GFP*). (**e**). Anti-GFP (green) immunostaining revealing Annexin-V binding to the longitudinal visceral muscles in unchallenged wild-type flies (*lpp>AnnexinV-GFP*). Blue, Chitin; Green, GFP; Scale bar: 20μm.

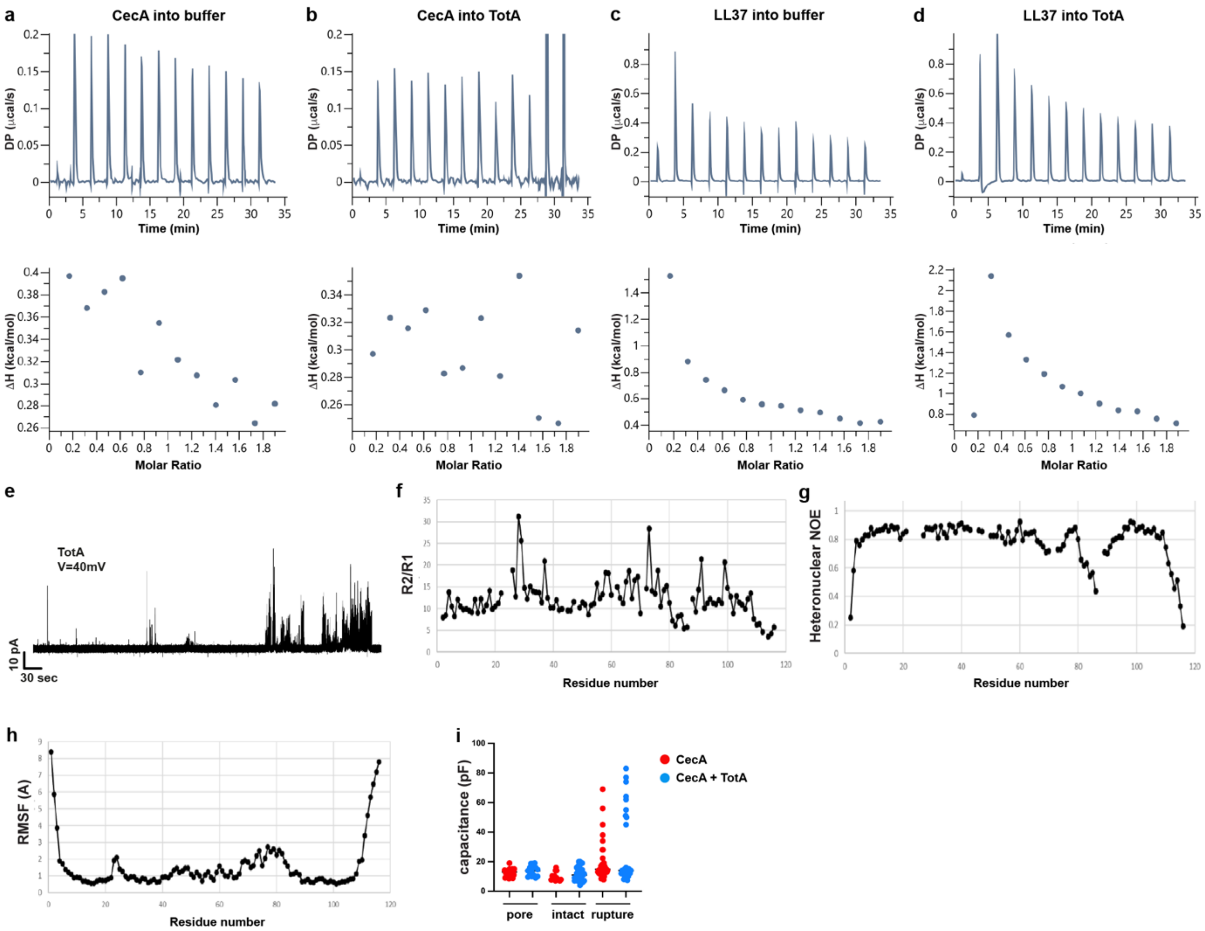

**Extended Data Figure 5: Isothermal titration calorimetry and TotA dynamics:** Isothermal titration calorimetry experiments reveal no binding of TotA to AMPs. Top panels show Representative raw thermogram plots of Isothermal titration calorimetry experiments. (**a,b**) 2 µL of 2mM CecA were injected 13 times in a chamber containing buffer alone (**a**) or 200 µM TotA (**b**) Lower panels show the integrated data versus molar ratio of peptide to TotA. (**c,d**) 2 µL of 2mM LL37 were injected 13 times in a chamber containing buffer alone (**c**) or 200 µM TotA (**d**). (**e**) Current recordings through lipid bilayer after addition of 4 µM TotA. (**f-h**) Results from $^{15}$N relaxation (**g**, $R_2/R_1$ ratio, **g**, heteronuclear NOE) compared to the flexibility observed in the NMR ensemble (**h**, RMSF to mean structure). In the $R_2/R_1$ plot, the median reflects a correlation time of 7.45 ns, perfectly consistent with a monomeric, globular, 12 kDa protein as TotA. Positive deviations of the ratio hint at residues experiencing slow exchange, and negative deviations hint at residues undergoing fast dynamics. Likewise, low heteronuclear NOE values indicate fast dynamics, and high RMSF points at flexibility and/or uncertainty in the NMR ensemble. **i**. Capacitances (in pF) of membrane bilayers upon addition of CecA (red) or CecA + TotA (blue) upon pore formation, membrane rupture or when the membrane stayed intact.

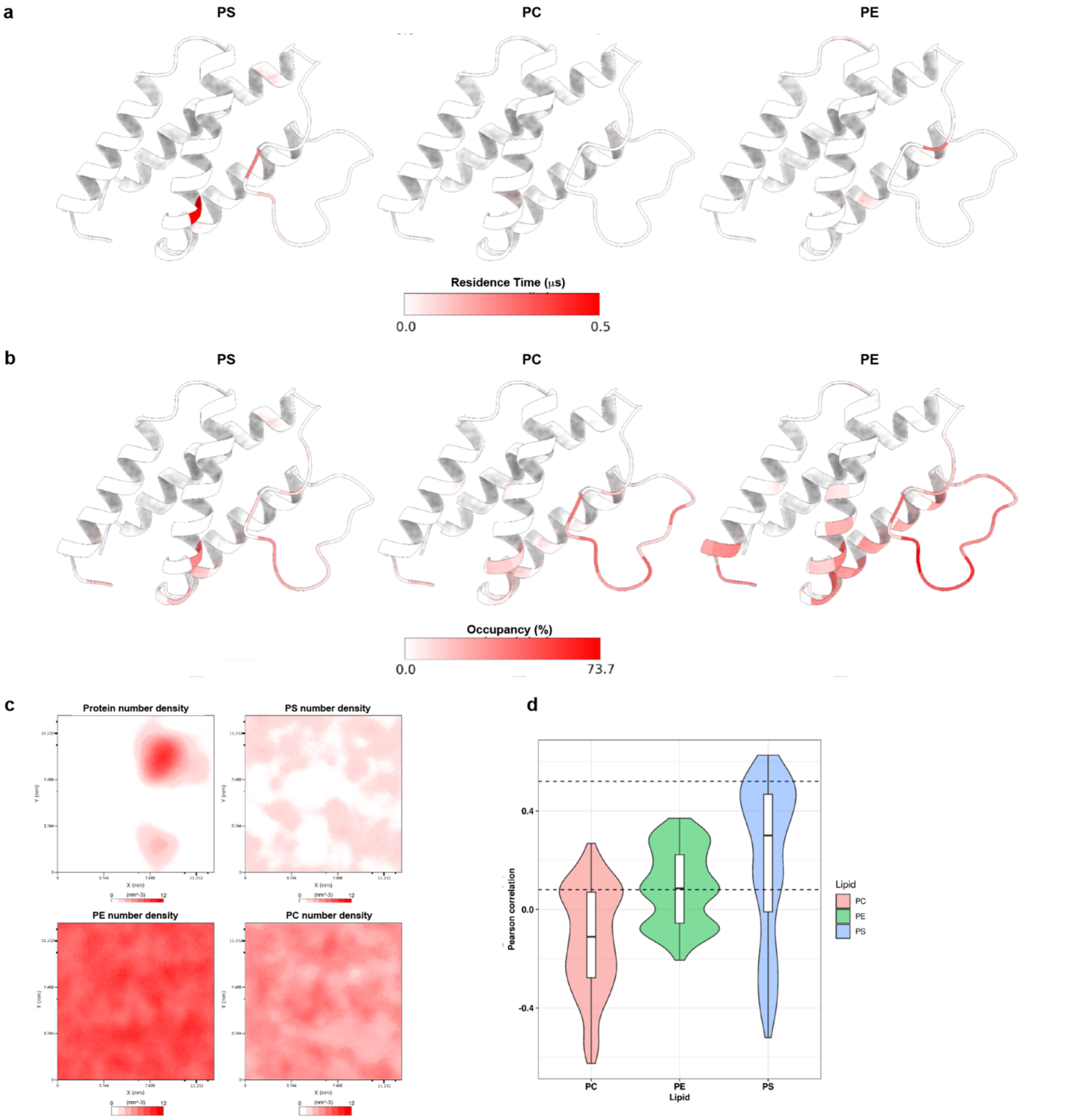

**Extended Data Figure 6: Molecular Dynamics (MD) simulations reveal specific binding of TotA to PS:** Residue-wise residence times (**a**) and occupancy (**b**) of TotA with PS, PC and PE during the MD simulations. Despite PS displaying the least occupancy as PS is less abundant that the two others (**b**), it does possess the highest residence times during the simulations (**a**). This suggests that TotA has a high specificity towards this lipid. Three TotA residues, R59, R95 and K96, contributed the most to PS specificity based on their residence times ($t^{R59}$ = 0.500 μs, $t^{R95}$ = 0.057 μs, and $t^{K96}$ = 0.154 μs) and occupancy (percentage of simulation time bound to PS, $Occup.^{R59}$ = 39.4%, $Occup.^{R95}$ = 33.6%, and $Occup.^{K96}$ = 36.0%). (**c-d**) TotA and lipid distributions on the XY plane viewed from above the membrane. (**c**) Number density plot of TotA (Protein) and the three lipids during the MD simulations. TotA clustered more to PS when compared to PC and PE. (**d**) Distribution of the correlation between the number density of TotA and the lipid. The position of TotA was more frequently correlated with that of the PS molecules then PC and PE. The dashed lines indicated the mode of the correlation distribution (0.52, for PS, 0.08 for PC and PE).

.

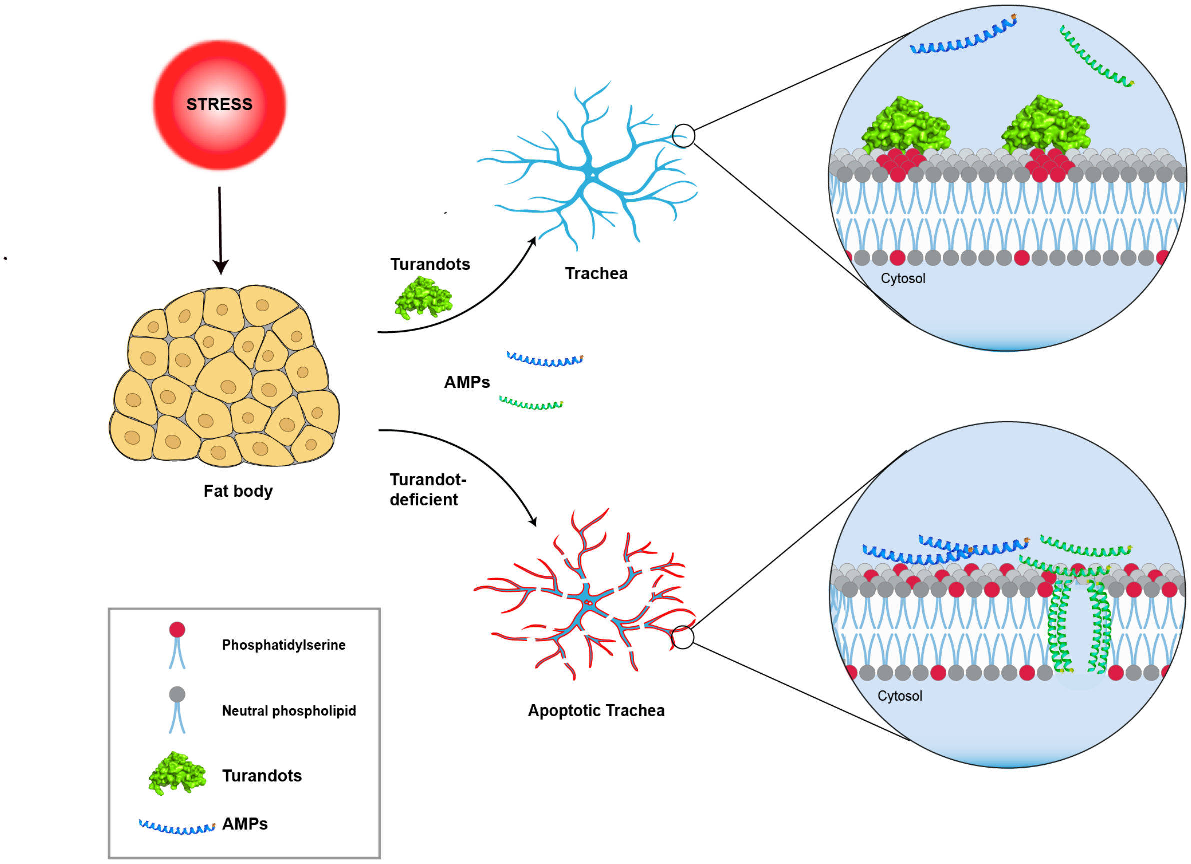

**Extended Data Figure 7: Turandots promote stress resilience by protecting *Drosophila* trachea from antimicrobial peptides.**
Certain tissues such as trachea expose unusually high amounts of phosphatidylserine even in basal conditions, sensitizing these tissues to cationic antimicrobial peptides. Upon stress exposure, Turandot proteins (green) are secreted into the hemolymph and bind to the membrane of tracheas. (Top) Turandot binding to phosphatidylserine (red) clusters this negatively charged lipid, inhibiting the formation of pores by AMPs (blue and light green), (Bottom) In absence of Turandots, cationic AMPs bind to trachea provoking tracheal apoptosis and lower resilience to stress.

**Extended Data Table 1: List of *Drosophila* stocks used in this study**

| figure | full genotype | short name | origin |
|---|---|---|---|
| **figure 1** | Iso $w^{1118}$ DrosDel | w iso | Ferreira et al. 2014 |
| | iso;iso;$TotAZ^{SK6}$ | TotAZ | this study |
| | iso;$totM^{JP78}$/CyO | TotM | this study |
| | iso;$totM^{JP1621}$;$TotX^{JP44}$ | TotMX | this study |
| | iso;$totM^{JP1621}$;$TotAZ^{SK6}$ | TotMAZ | this study |
| | iso;iso;$TotX^{JP44}$, $TotAZ^{SK6}$ | TotAZX | this study |
| | iso;$totM^{JP1621}$;$TotX^{JP44}$, TotAZ $^{SK6}$ | TotXMAZ | this study |
| | iso;iso;$totX^{JP147}$ | totX | this study |
| | iso;$totM^{JP1621}$ ;iso | TotM | this study |
| | $w^{1118}$;$totM^{JP1621}$/CyO; lpp-gal4,$TotX^{JP44}$, $TotAZ^{SK6}$/TM3sb | lpp XMAZ | this study |
| | ;UAS-TotA-HA @attP1, $totM^{JP1621}$/CyO; $TotAZ^{SK6}$, $totX^{JP44}$ | UAS-totA XMAZ | this study |
| **figure 2** | iso $w^{1118}$ DrosDel | w iso | Ferreira et al. 2014 |
| | w;tub-$gal80^{ts}$;lpp-gal4 | Lpp> | Brankatschk and Eaton, 2010 |
| | ;UAS-HA5UTR-TotA @attP40 | UAS-HA-TotA | this study |
| | iso;$totM^{JP1621}$;$TotX^{JP44}$, $TotAZ^{SK6}$ | TotXMAZ | this study |
| | $w^{1118}$;P(GD6210)v14415 | TotA RNAi | VDRC 14415 |
| **figure 3** | iso $w^{1118}$ DrosDel | w iso | Ferreira et al. 2014 |
| | iso;$totM^{JP1621}$;$TotX^{JP44}$, TotAZ $^{SK6}$ | TotXMAZ | this study |
| | $w^{1118}$;$totM^{JP1621}$/CyO; lpp-gal4,$TotX^{JP44}$, TotAZ $^{SK6}$/TM3sb | lpp XMAZ | this study |
| | ;UAS-TotA-HA @attP1, $totM^{JP1621}$/CyO; TotAZ $^{SK6}$, $totX^{JP44}$ | UAS-totA XMAZ | this study |
| | ; ; dsrf-gal4, UAS-PH-GFP, $TotX^{JP44}$, TotAZ $^{SK6}$ | dsrf>GFP AZX | Gervais and Casanova 2010 and this study |
| | $w^{1118}$;; hsp-nls-Timer | timer | Lidsky et al. 2018 |
| | ;; hsp-nls-Timer, $TotX^{JP44}$, $TotAZ^{SK6}$ | timer AZX | this study |
| | ;btl-gal4,UAS-GFP; tub-$gal80^{ts}$ | btl(ts)>GFP | BL8807 and this study |
| | ;btl-gal4, UAS-GFP;TotAZ $^{SK6}$, $totX^{JP44,}$ tub-$gal80^{ts}$ | btl(ts)>GFP AZX | this study |
| | w[*]; P{w[+mC]=UAS-p35.H}BH1,TotXJP44, $TotAZ^{SK6}$ | p35 AZX | BL5072 and this study |
| **Figure 4** | ;;UAS-sec-GFP | sec-GFP | Fabrowski et al. 2013 and this study |
| | ;;UAS-sec-GFP,TotAZ $^{SK6}$, $totX^{JP44}$ | sec-GFP AZX | Fabrowski et al. 2013 and this study |
| | ;UAS-AnnexinV-GFP | AnnexinV-GFP | Sapar et al. 2018 |
| | ;UAS-AnnexinV-GFP;$TotAZ^{SK6}$, $totX^{JP44}$ | AnnexinV-GFP AZX | Sapar et al. 2018 and this study |
| | ;btl-gal4, UAS-GFP;$TotAZ^{SK6}$, $totX^{JP44}$, tub-$gal80^{(ts)}$ | btl(ts)>GFP AZX | this study |
| | ;scramb1 IR,TotAZ $^{SK6}$, $totX^{JP44}$ | scramb1 IR AZX | v107024 and this study |
| | ;btl-gal4, UAS-GFP; tub-$gal80^{(ts)}$ | btl(ts)>GFP | BL8807 and this study |
| | iso;iso;$Rel^{E20}$ | RelE20 | BL55714 |
| | $Def^{SK3}$, $AttC^{Mi}$, Dro-$AttAB^{SK2}$, $Mtk^{R1}$, $Dpt^{SK1}$; $Drs^{R1}$, CecKOΔA-C, $AttD^{SK1}$ | AMP14 | Carboni et al. 2022 |
| | ;UAS-xkr8/CyoGFP | UAS-xkr8 | Sapar et al. 2018 |
| | ;btl-gal4, UAS-GFP; | btl>GFP | BL8807 |
| | ;btl-gal4, UAS-GFP; $CecKO^{\Delta A\text{-}C}$ | btl>GFP; CecAC | BL8807 and this study |
| | ;UAS-xkr8/CyoGFP; $CecKO^{\Delta A\text{-}C}$ | UAS-xkr8; CecAC | Sapar et al. 2018 and this study |
| | iso;DefSK3, $AttC^{Mi}$, Dro-$AttAB^{SK2}$, $Mtk^{R1}$, $Dpt^{SK1}$; iso;iso;$CecKO^{\Delta A\text{-}C}$,$TotX^{JP44}$, $TotAZ^{SK6}$ | AMP12,AZX | this study |
| | iso $w^{1118}$ DrosDel | w iso | Ferreira et al. 2014 |
| | iso;iso;$TotX^{JP44}$, $TotAZ^{SK6}$ | TotAZX | this study |
| | $w^{1118}$;; lpp-gal4,$TotX^{JP44}$, $TotAZ^{SK6}$/TM3sb | lpp> AZX | this study |
| **Extended data figure 1** | $w^{1118}$;$totM^{JP1621}$, act-gal4/CyO; $TotX^{JP44}$, $TotAZ^{SK6}$/TM3sb | act> XMAZ | BL3853 and this study |
| | iso $w^{1118}$ DrosDel | w iso | Ferreira et al. 2014 |
| | ;UAS-TotA-HA @attP1, $totM^{JP1621}$/CyO; TotAZ, $totX^{JP44}$ | UAS-totA XMAZ | this study |
| | iso;$totM^{JP1621}$;$TotX^{JP44}$, TotAZ $^{SK6}$ | TotXMAZ | this study |
| **Extended data figure 2** | w;tub-$gal80^{ts}$;lpp-gal4 | Lpp> | Brankatschk and Eaton, 2010 |
| | ;UAS-TotM-HA@attP1 | TotM-HA | this study |
| | iso $w^{1118}$ DrosDel | w iso | Ferreira et al. 2014 |
| | ;UAS-TotA-HA @attP1 | TotA-HA | this study |
| **Extended data figure 3** | iso $w^{1118}$ DrosDel | w iso | Ferreira et al. 2014 |
| | iso;$totM^{JP1621}$;$TotX^{JP44}$, TotAZ $^{SK6}$ | TotXMAZ | this study |
| **Extended data figure 4** | $w^{1118}$;tub-$gal80^{ts}$;lpp-gal4 | Lpp> | Brankatschk and Eaton, 2010 |
| | $w^{1118}$;; lpp-gal4,$TotX^{JP44}$, $TotAZ^{SK6}$/TM3sb | lpp> AZX | this study |
| | ;UAS-AnnexinV-GFP | AnnexinV-GFP | Sapar et al. 2018 |
| | ;UAS-AnnexinV-GFP;$TotAZ^{SK6}$, $totX^{JP44}$ | UAS-AnnexinV-GFP;AZX | Sapar et al. 2018 and this study |

| | | |
|---|---|---|
| $w^{1118}$;UAS-GFP-LactC1C2 | UAS-lactGFP | Sapar et al. 2018 |

### Extended Data Table 2: NMR and refinement statistics for protein structures

| | Protein |
|---|---|
| **NMR distance and dihedral constraints** | |
| Distance constraints | |
| Total NOE | 1061 |
| Intra-residue | 225 |
| Inter-residue | 836 |
| Sequential ($\|i - j\| = 1$) | 352 |
| Medium-range ($\|i - j\| < 4$) | 282 |
| Long-range ($\|i - j\| > 5$) | 140 |
| Intermolecular | -- |
| Hydrogen bonds | -- |
| Total dihedral angle restraints (from Talos+) | 140 |
| $\phi$ | 70 |
| $\psi$ | 70 |
| **Structure statistics** | |
| Violations (mean and s.d.) | |
| Distance constraints (Å) | 0.5 ± 0.3 |
| Dihedral angle constraints (º) | 7.5 ± 2.6 |
| Max. dihedral angle violation (º) | 25 |
| Max. distance constraint violation (Å) | 1.47 |
| Deviations from idealized geometry | |
| Bond lengths (Å) | 0 |
| Bond angles (º) | 0 |
| Impropers (º) | 0 |
| Average pairwise r.m.s. deviation* (Å) | |
| Heavy | 3.97 ± 2.37 |
| Backbone | 3.28 ± 1.31 |

* Pairwise r.m.s. deviation was calculated among 20 refined structures throughout the full protein lengths.

### Extended Data Table 3: Statistics

| | | | n | test | tailed | correction | group1 | group2 | pvalue | pvalue_bonferroni |
|---|---|---|---|---|---|---|---|---|---|---|
| | A | w | 120 | coxPH | N/A | N/A | | | $10^{-5}$ | |
| | | XMAZ | 120 | coxPH | N/A | N/A | | | | |
| | | w | 120 | coxPH | N/A | N/A | | | $10^{-16}$ | |
| | B | XMAZ | 121 | coxPH | N/A | N/A | | | | |
| | C | w | 120 | coxPH | N/A | N/A | | | $10^{-16}$ | |
| | | XMAZ | 120 | coxPH | N/A | N/A | | | | |
| | | UAS-totA;XMAZ | 60 | coxPH | N/A | bonferonni | | | | |
| figure 1 | | lpp>+;XMAZ | 80 | coxPH | N/A | bonferonni | | | | |
| | | lpp>totA;XMAZ | 80 | coxPH | N/A | bonferonni | | | | |
| | | | | | | | UAS-totA;XMAZ | lpp>+;XMAZ | 1.19341586034287e-08 | 7.16049516205724e-08 |
| | E | | | | | | UAS-totA;XMAZ | lpp>TotA;XMAZ | 1.45061994768379e-18 | 8.70371968610274e-18 |
| | | multiple comparisons | | | | | lpp>+;XMAZ | UAS-totA;XMAZ | 1.19341586034287e-08 | 7.16049516205724e-08 |
| | | | | | | | lpp>+;XMAZ | lpp>TotA;XMAZ | 2.92399262162095e-06 | 1.75439557297257e-05 |
| | | | | | | | lpp>TotA;XMAZ | UAS-totA;XMAZ | 1.45061994768379e-18 | 8.70371968610274e-18 |
| | | | | | | | lpp>TotA;XMAZ | lpp>+;XMAZ | 2.92399262162095e-06 | 1.75439557297257e-05 |
| | | w | 24 | one way ANOVA | one-tailed | dunnett's | | | | |
| | B | XMAZ | 26 | one way ANOVA | one-tailed | dunnett's | | | <0.0001 | |
| | | w nacl | 33 | one way ANOVA | one-tailed | dunnett's | | | <0.0001 | |
| | | XMAZ nacl | 31 | one way ANOVA | one-tailed | dunnett's | | | <0.0001 | |
| | | w | 26 | one way ANOVA | one-tailed | dunnett's | | | | |
| | C | XMAZ | 29 | one way ANOVA | one-tailed | dunnett's | | | 0,0032 | |
| | | w nacl | 34 | one way ANOVA | one-tailed | dunnett's | | | 0,06 | |
| | | XMAZ nacl | 31 | one way ANOVA | one-tailed | dunnett's | | | 0,0002 | |
| | | lpp>w XMAZ+/- | 22 | one way ANOVA | one-tailed | dunnett's | | | | |
| | E | lpp>w XMAZ-/- | 21 | one way ANOVA | one-tailed | dunnett's | | | 0,0005 | |
| | | lpp>TotA-HA XMAZ-/- | 22 | one way ANOVA | one-tailed | dunnett's | | | 0,2873 | |
| | | UAS-TotA-HA XMAZ-/- | 7 | one way ANOVA | one-tailed | dunnett's | | | 0,009 | |
| | F | hs>nls-timer AZX+/- | 12 | mann-whitney | two-tailed | N/A | | | <0.0001 | |
| | | hs>nls-timer AZX-/- | 9 | mann-whitney | two-tailed | N/A | | | <0.0001 | |
| | G | w | 408 | Welch two sample t-test | | N/A | | | <0.05 | |
| | | XMAZ | 303 | Welch two sample t-test | | N/A | | | <0.05 | |
| | | w normoxia | 49 | coxPH | N/A | bonferonni | | | | |
| | | XMAZ normoxia | 80 | coxPH | N/A | bonferonni | | | | |
| | | w hypoxia | 77 | coxPH | N/A | bonferonni | | | | |
| | | XMAZ hypoxia | 77 | coxPH | N/A | bonferonni | | | | |
| | H | | | | | | w normoxia | XMAZ normoxia | 2,34E-15 | 2,81E-14 |
| | | | | | | | w normoxia | w hypoxia (4% O2) | 6,84E-10 | 8,21E-09 |
| | | multiple comparisons | | | | | w normoxia | XMAZ hypoxia (4% O2 | 7,14E-19 | 8,57E-18 |
| | | | | | | | XMAZ normoxia | w hypoxia (4% O2) | 0,000118997 | 0,001427969 |
| | | | | | | | XMAZ normoxia | XMAZ hypoxia (4% O2 | 3,06E-10 | 3,68E-09 |
| | | | | | | | w hypoxia (4% O2) | XMAZ hypoxia (4% O2 | 1,01E-17 | 1,21E-16 |
| figure 3 | | w normoxia | 162 | coxPH | N/A | bonferonni | | | | |
| | | XMAZ normoxia | 162 | coxPH | N/A | bonferonni | | | | |
| | | w hyperoxia | 160 | coxPH | N/A | bonferonni | | | | |
| | | XMAZ hyperoxia | 155 | coxPH | N/A | bonferonni | | | | |
| | I | | | | | | w normoxia | XMAZ | 5.8220943289877e-37 | 6.98651319478523e-36 |
| | | | | | | | w normoxia | w hyperoxia (40% O2) | 0.858922806634268 | 1 |
| | | multiple comparisons | | | | | w normoxia | XMAZ hyperoxia (40% | 1.17609999418142e-23 | 1.41131999301771e-22 |
| | | | | | | | XMAZ | w hyperoxia (40% O2) | 1.78262870005903e-32 | 2.13915444007084e-31 |
| | | | | | | | XMAZ | XMAZ hyperoxia (40% | 2.6441909896877e-06 | 3.17302918762524e-05 |
| | | | | | | | w hyperoxia (40% O | XMAZ hyperoxia (40% | 2.0359564217623e-19 | 2.44314770611476e-18 |
| | K | w | 21 | mann-whitney | two-tailed | N/A | | | <0.0001 | |
| | | XMAZ | 20 | mann-whitney | two-tailed | N/A | | | | |
| | | btl(ts)>w AZX +/- ct | 15 | one way ANOVA | one-tailed | dunnett's | | | | |
| | M | btl(ts)>w AZX -/- ct | 14 | one way ANOVA | one-tailed | dunnett's | | | 0,0002 | |
| | | UAS p35 AZX -/- ct | 11 | one way ANOVA | one-tailed | dunnett's | | | <0.0001 | |
| | | btl(ts)>p35 AZX -/- ct | 11 | one way ANOVA | one-tailed | dunnett's | | | 0,8899 | |
| | | btl(ts)>w AZX +/- ct | 58 | coxPH | N/A | bonferonni | | | | |
| | | btl(ts)>w AZX -/- ct | 44 | coxPH | N/A | bonferonni | | | | |
| | | UAS p35 AZX -/- ct | 58 | coxPH | N/A | bonferonni | | | | |
| | | btl(ts)>p35 AZX -/- ct | 62 | coxPH | N/A | bonferonni | | | | |
| | N | | | | | | control | btl(ts)>w AZX -/- | 0.00155122498997267 | 0.018614699879672 |
| | | | | | | | control | btl(ts)>p35 AZX -/- | 0.317688812456521 | 1 |
| | | multiple comparisons | | | | | control | UAS p35 AZX -/- | 5.23839801496303e-10 | 6.28607761795564e-09 |
| | | | | | | | btl(ts)>w AZX -/- | btl(ts)>p35 AZX -/- | 2.80956090437762e-05 | 0.000337147308525315 |
| | | | | | | | btl(ts)>w AZX -/- | UAS p35 AZX -/- | 0.00238821234797067 | 0.0286585481756481 |
| | | | | | | | btl(ts)>p35 AZX -/- | UAS p35 AZX -/- | 4.69905391510161e-13 | 5.63886469812194e-12 |
| | | btl(ts)>w AZX +/- | 21 | one way ANOVA | one-tailed | dunnett's | | | | |
| | B | btl(ts)>w AZX -/- | 22 | one way ANOVA | one-tailed | dunnett's | | | <0.0001 | |
| | | +/scramb1 IR AZX -/- | 20 | one way ANOVA | one-tailed | dunnett's | | | <0.0001 | |
| | | btl(ts)> scramb1 IR AZX -/- | 21 | one way ANOVA | one-tailed | dunnett's | | | 0,0231 | |
| | | btl(ts)>w AZX +/- | 101 | coxPH | N/A | bonferonni | | | | |
| | | btl(ts)>w AZX -/- | 98 | coxPH | N/A | bonferonni | | | | |
| | | +/scramb1 IR AZX -/- | 99 | coxPH | N/A | bonferonni | | | | |
| | | btl(ts)> scramb1 IR AZX -/- | 100 | coxPH | N/A | bonferonni | | | | |
| | C | | | | | | +/scramb1 IR;AZX -/ | btl(ts)>w AZX +/- | 3.65123058756842e-16 | 1.09536917627052e-14 |
| | | | | | | | +/scramb1 IR;AZX -/ | btl(ts)>w AZX -/- | 2.22769354595678e-06 | 6.68308063787034e-05 |
| | | multiple comparisons | | | | | +/scramb1 IR;AZX -/ | btl(ts)>scramb1 IR;AZ) | 1.00358027556739e-24 | 3.01074082670218e-23 |
| | | | | | | | btl(ts)>w AZX +/- | btl(ts)>w AZX -/- | 1.8454783536133e-05 | 0.00055364350608399 |
| | | | | | | | btl(ts)>w AZX +/- | btl(ts)>scramb1 IR;AZ) | 0.676638942442102 | 1 |
| | | | | | | | btl(ts)>scramb1 IR;A | btl(ts)>w AZX -/- | 1.52113068748423e-08 | 4.56339206245268e-07 |
| | D | btl(ts)>GFP, w | 22 | mann-whitney | two-tailed | N/A | | | <0.0001 | |
| | | btl(ts)>GFP, xkr8 | 22 | mann-whitney | two-tailed | N/A | | | | |
| | E | btl(ts)>GFP, w | 130 | coxPH | N/A | N/A | | | 6.19887959845118e-14 | |
| | | btl(ts)>GFP, xkr8 | 183 | coxPH | N/A | N/A | | | | |
| | | w ct | 32 | one way ANOVA | one-tailed | dunnett's | | | | |
| | | w HK | 31 | one way ANOVA | one-tailed | dunnett's | | | | <0.0001 |
| | F | rel | 23 | one way ANOVA | one-tailed | dunnett's | | | | 0,9916 |
| | | rel HK | 23 | one way ANOVA | one-tailed | dunnett's | | | | 0,9934 |
| | | AMP14 ct | 16 | one way ANOVA | one-tailed | dunnett's | | | | 0,9997 |
| Figure 4 | | AMP14 HK | 16 | one way ANOVA | one-tailed | dunnett's | | | | 0,9994 |
| | | btl>GFP, w | 36 | one way ANOVA | one-tailed | dunnett's | | | | |
| | G | btl>GFP, xkr8 | 32 | one way ANOVA | one-tailed | dunnett's | | | | <0.0001 |
| | | btl>GFP, w cecAC | 20 | one way ANOVA | one-tailed | dunnett's | | | | 0,1856 |
| | | btl>GFP, xkr8 cecAC | 21 | one way ANOVA | one-tailed | dunnett's | | | | 0,9983 |
| | | btl>GFP, w | 95 | coxPH | N/A | bonferonni | | | | |
| | | btl>GFP, xkr8 | 103 | coxPH | N/A | bonferonni | | | | |
| | | btl>GFP, w cecAC | 129 | coxPH | N/A | bonferonni | | | | |
| | | btl>GFP, xkr8 cecAC | 129 | coxPH | N/A | bonferonni | | | | |
| | H | | | | | | btl>w | btl>xkr8 | 4.44221922561541e-06 | 0.000186573207475847 |
| | | | | | | | btl>w | btl >w CecAC -/- | 1.15191745813471e-06 | 4.83805332416579e-05 |
| | | multiple comparisons | | | | | btl>w | btl >xkr8 CecAC -/- | 0.757618830871803 | 1 |
| | | | | | | | btl>xkr8 | btl >w CecAC -/- | 1.40810770537736e-20 | 5.91405236258491e-19 |
| | | | | | | | btl>xkr8 | btl >xkr8 CecAC -/- | 1.28256149937143e-07 | 5.38675829736001e-06 |
| | | | | | | | btl >w CecAC -/- | btl >xkr8 CecAC -/- | 7.17595891903314e-10 | 3.01390274599392e-08 |
| | | w | 23 | one way ANOVA | one-tailed | dunnett's | | | | |
| | I | AZX-/- | 20 | one way ANOVA | one-tailed | dunnett's | | | | 0,0005 |
| | | AMP2; CecAC, TotAZX | 19 | one way ANOVA | one-tailed | dunnett's | | | | 0,1776 |
| | | w | 79 | coxPH | N/A | bonferonni | | | | |
| | | AZX-/- | 121 | coxPH | N/A | bonferonni | | | | |
| | J | AMP2; CecAC, TotAZX | 118 | coxPH | N/A | bonferonni | | | | |
| | | | | | | | w | TotAZX | 5.42521802542149e-17 | 3.25513081525289e-16 |
| | | multiple comparisons | | | | | w | AMP2; CecAC, TotAZX | 0.281449057946893 | 1 |
| | | | | | | | TotAZX | AMP2; CecAC, TotAZX | 3.77152156397125e-15 | 2.26291293838275e-14 |
| | | ctrl | 9 | one way ANOVA | one-tailed | Holm-Šídák's | | | | |
| | A | LL37 | 3 | one way ANOVA | one-tailed | Holm-Šídák's | | | | <0.0001 |
| | | LL37 + TotA | 3 | one way ANOVA | one-tailed | Holm-Šídák's | | | | <0.0001 |
| Figure 5 | | ctrl | 9 | one way ANOVA | one-tailed | Holm-Šídák's | | | | |
| | B | LL37 | 9 | one way ANOVA | one-tailed | Holm-Šídák's | | | | <0.0001 |
| | | LL37 + TotA | 9 | one way ANOVA | one-tailed | Holm-Šídák's | | | | 0,0415 |
